# Chronic Polystyrene Nanoplastics Exposure Reprograms Gene Expression, Alternative Splicing, and Disrupts Host–Microbiome–Metabolic Networks to Promote Atherosclerosis in LDLr⁻/⁻ Mice

**DOI:** 10.64898/2026.07.09.736446

**Authors:** Ajmal Khan, Delicia Esther Cardenas Vasquez, Yaru Si, Warren S. Vidar, Kerui Wu, Norman Chiu, Zhenquan Jia

**Affiliations:** Department of Biology, University of North Carolina at Greensboro, Greensboro, NC 27412, USA; Joint School of Nanoscience and Nanoengineering, University of North Carolina at Greensboro, Greensboro, NC 27412, USA; Department of Chemistry and Biochemistry, University of North Carolina at Greensboro, Greensboro, NC 27412, USA

**Keywords:** Nanoplastics, Atherosclerosis, Transcriptomics and Alternative splicing, Metagenomics, Metabolomics

## Abstract

Although micro- and nanoplastics have been detected in human atherosclerotic plaques, their mechanistic contribution to disease pathogenesis remains poorly defined. Most experimental studies have used microplastics (particles > 1 μm) in non-atherosclerotic animal models or the ApoE⁻/⁻ mouse, relying on short-term exposure or single-pathway analyses, whereas the chronic cardiovascular effects of nanoplastics (< 100 nm) remain exceedingly scarce—despite their higher biological reactivity and greater tissue penetrance. To address this gap, this study employs a multi-omics approach to investigate the chronic (12-week) oral exposure to 80 nm polystyrene nanoplastics in LDLr⁻/⁻ mice. We uniquely integrate aortic plaque quantification, hepatic transcriptomics with global alternative splicing profiling, gut microbiome 16S sequencing, and liver untargeted metabolomics to construct a unified host–microbiome–metabolite network. Nanoplastic exposure significantly exacerbates aortic lipid deposition, suppresses hepatic detoxification and anti-atherogenic lipid pathways primarily through transcriptional and post-transcriptional level changes driven by alternative splicing events (e.g., intron retention and isoform switching), and induces gut dysbiosis marked by a reduction in SCFA-producing commensals and enrichment of pro-atherogenic pathobionts—perturbations that correlate with specific hepatic functional modules. Metabolomic changes, including decreased levels of the glutathione precursor γ-glutamylcysteine and the choline-derived metabolite neurine, implicate oxidative stress and TMAO-related pathways. Cross-species validation using human atherosclerotic transcriptomic and metagenomic datasets supports the clinical translatability. By integrating multi-level biological responses, this work establishes nanoplastics as an environmental cardiovascular risk factor and uncovers novel regulatory mechanisms involving splicing-associated transcriptional reprogramming and gut–liver crosstalk, offering potential early-warning biomarkers and therapeutic targets for nanoplastic-associated cardiovascular disease.

**Highlights:**

- Exposure to polystyrene nanoplastics (80 nm) increases aortic lipid burden in LDLr⁻/⁻ mice.
- Alternative splicing and isoform switching were identified as novel hepatic responses.
- SCFA-producing gut commensals are depleted, and pathobionts are enriched in response to nanoplastics.
- A gut–liver network links suppressed detoxification to gut microbial dysbiosis.
- Mouse transcriptomics and metagenomics overlap with human atherosclerosis omics datasets.

Graphical Abstract

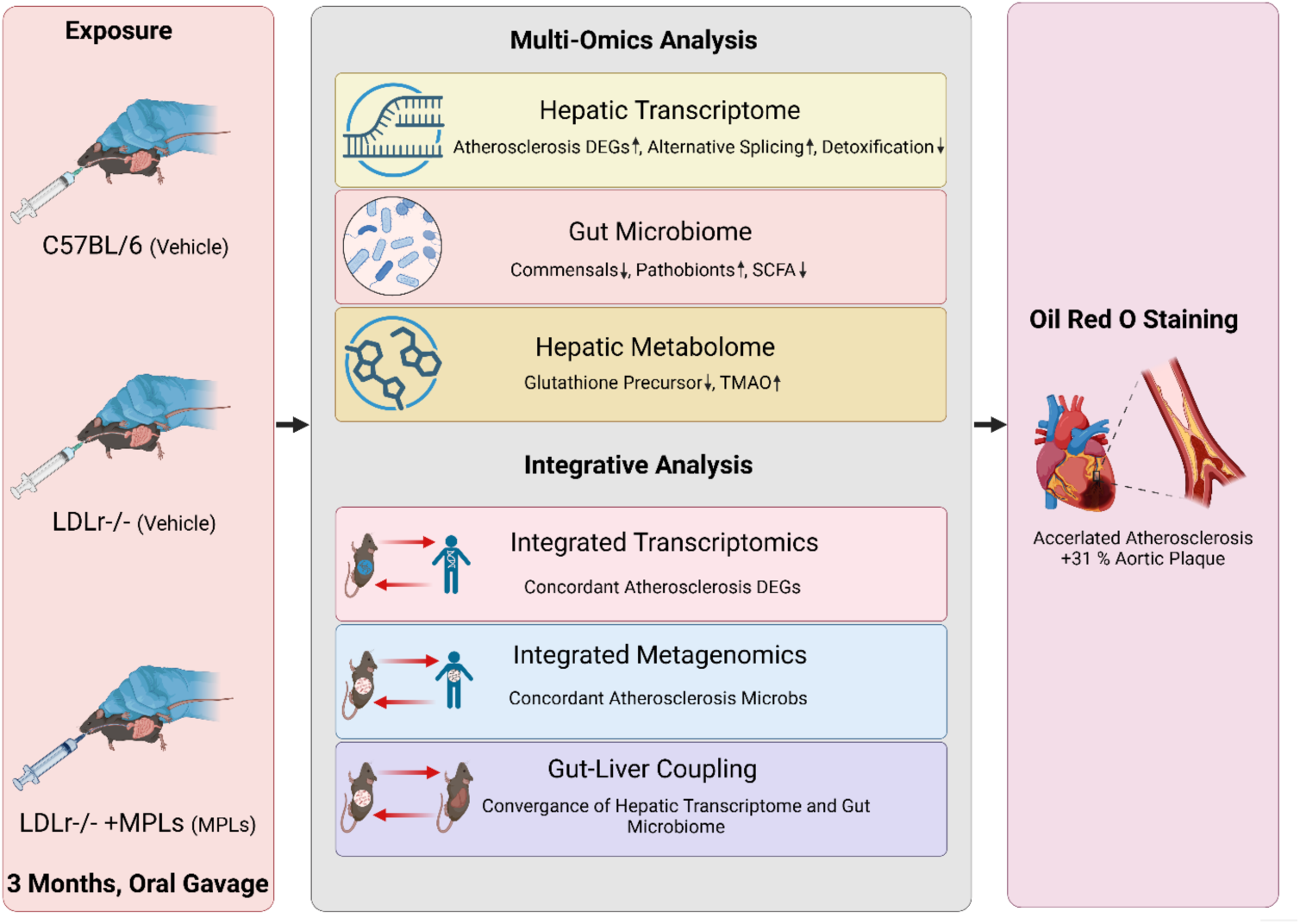

## Introduction

Cardiovascular diseases (CVDs) are responsible for approximately 20.5 million deaths each year (roughly one-third of all global deaths). Hence, they are touted as the number one killer disease (Federation 2023). Atherosclerosis, the chronic inflammatory form of CVDs, is characterized by the thickening of blood vessels caused by the build-up of plaque (deposits of fatty substances, cholesterol, cellular waste products, calcium, and fibrin) in the inner lining of blood vessels (Hansson 2005, Ait-Oufella, Taleb et al. 2011). Despite the multifactorial nature of atherosclerosis pathogenesis, growing evidence implicates environmental contaminants, such as smoke, chemicals, and particulate pollutants, as contributing factors in atherogenesis (Bhatnagar 2006, Cosselman, Navas-Acien et al. 2015, Shukla, Chitrakar et al. 2020). Among these, micro- and nanoplastics, which are present in the human food chain, drinking water, and air (Khan and Jia 2023) have been detected in various human tissues, including brain, placenta, lungs, kidneys, blood, and cardiovascular tissues (Schwabl, Koppel et al. 2019, Ragusa, Svelato et al. 2021, Huang, Huang et al. 2022, Jenner, Rotchell et al. 2022, Leslie, van Velzen et al. 2022), are linked to various human diseases such as neurodegenerative diseases (Weiss and Ding 2024), and ovarian fibrosis (An, Wang et al. 2021). Most importantly for cardiovascular risk, micro- and nanoplastics have been detected in human carotid atheromas, where their presence predicts myocardial infarction and stroke (Marfella, Prattichizzo et al. 2024). Experimental studies have linked microplastic and nanoplastics exposure to cardiac fibrosis, endothelial dysfunction, and myocardial damage (Hu and Palić 2020, Li, Zhu et al. 2020, Chen, Chen et al. 2023). However, the mechanism by which nanoplastics promote atherosclerosis remains unclear.

Experimental and epidemiological studies have shown that micro- and nanoplastics can disrupt the integrity of the intestinal barrier (Zeng, Li et al. 2024), induce microbial dysbiosis (Qiao, Sheng et al. 2019, Procopio, Soggiu et al. 2025), and cause systemic inflammation (Qiao, Sheng et al. 2019) and metabolic perturbations (Ye, Zhang et al. 2021). These effects overlap with key pathways and processes already known to be involved in the progression of atherosclerosis, such as immune activation, oxidative stress, and lipid dysregulation (Liu, Wang et al. 2023). Studies have further linked polystyrene nanoplastics to artery stiffening, plaque formation, macrophage M1 activation via MARCO upregulation, and acylcarnitine accumulation in ApoE⁻/⁻ mice (Wang, Liang et al. 2023). Consistent with this, oral exposure to polystyrene nanoplastics promotes aortic plaque formation and disrupts hepatic lipid metabolism in ApoE⁻/⁻ mice (Wen, Sun et al. 2024), while dietary polystyrene microplastics increase cardiac lipid accumulation and impair glucose homeostasis (Zhao, Gomes et al. 2024). Oral administration of 1 μm polystyrene microplastics accelerates the progression of atherosclerosis in LDLr⁻/⁻ mice in a sex-dependent manner (Lin, Pan et al. 2025). These studies establish an evidentiary foundation for micro- and nanoplastics as an environmental risk factor for cardiovascular disease. Nevertheless, the current body of knowledge suffers from several critical limitations that hinder a comprehensive understanding of microplastic cardiovascular toxicity. Most existing studies have relied on short-term exposure paradigms and non-atherosclerotic animal models, thereby providing only limited mechanistic insight into how chronic microplastic and nanoplastics exposure contributes to atherosclerosis development. Although accumulating evidence suggests that exposure to micro- and nanoplastics accelerates atherosclerotic progression in genetically susceptible hosts, the underlying mechanisms remain incompletely understood. Furthermore, the vast majority of investigations have been conducted in ApoE⁻/⁻ mice, while studies using LDLr⁻/⁻ mice remain scarce, despite their closer resemblance to the metabolic and lipoprotein abnormalities observed in human atherosclerosis. In addition, the role of particle size has received insufficient attention; specifically, the long-term cardiovascular effects of nanoplastics (<100 nm) remain largely unexplored, even though particle size is a critical determinant of biodistribution, cellular uptake, and toxicological response. Most importantly, existing studies have rarely moved beyond a single-pathway perspective and have largely failed to integrate transcriptional reprogramming, post-transcriptional splicing regulation, gut microbiome alterations, and systemic metabolic perturbations into a unified mechanistic framework. Given the established role of gut microbiota and their metabolites in the development of atherosclerosis, and considering that orally ingested nanoplastics interact with the intestinal microenvironment before entering the systemic circulation, employing a systems-level multi-omics approach is crucial for elucidating the complex biological pathways linking chronic nanoplastic exposure to cardiovascular disease.

To address these gaps, this study utilized LDLr⁻/⁻ mice as a model to systematically evaluate the pro-atherosclerotic effects of long-term (12-week) oral exposure to 80 nm polystyrene nanoplastics. By integrating aortic plaque quantification, hepatic transcriptomic sequencing, genome-wide alternative splicing analysis, gut microbiota 16S sequencing, and hepatic untargeted metabolomics, we comprehensively characterized the multi-level biological perturbations induced by nanoplastics. To the best of our knowledge, this is the first study to employ a system-level multi-omics strategy to systematically investigate the cardiovascular toxicity of polystyrene nanoplastics in an LDLr⁻/⁻ mouse model. Our findings not only reveal novel mechanisms by which long-term nanoplastic exposure drives atherosclerosis but also provide crucial clues for identifying the key molecular pathways linking environmental plastic pollution to cardiovascular disease risk.

## Materials and Methods

### 2.1. Study design and treatment

6-week-old male LDL receptor knockout mice (LDLr⁻/⁻) purchased from Jackson Laboratory, Bar Harbor, ME, acclimated for one week, were housed 3 per cage under standard animal experimental conditions (temperature, humidity, and CO2-controlled facility with 12 h light/dark) with ad libitum access to water and food. All LDLr⁻/⁻ animals were fed an atherogenic Teklad diet (TD.88137 adjusted-calories diet; 21.2% fat by weight, 42% kcal from fat, 0.2% total cholesterol, and 34% sucrose by weight) and divided randomly into the polystyrene nanoplastics group (LDLr⁻/⁻+MPLs, n = 9) and LDLr⁻/⁻ knockout control group (LDLr⁻/⁻, n = 9). Additional normal mice were used as normal controls (C57BL/6, n = 9 and were fed a normal diet. For the polystyrene nanoplastics group (LDLr⁻/⁻+MPLs), polystyrene nanoplastics (sulfate-stabilized linear polystyrene, average diameter 80 nm; 5.0% w/v; Cat# PP-008-100, Spherotech, Lake Forest, IL) at a dose of 200 mg/kg body weight, and for the control groups (LDLr⁻/⁻ and C57BL/6), an equivalent volume of vehicle was administered through oral gavage every other day for 3 months. Because the polystyrene nanoplastic stock is supplied in deionized water containing 0.02% sodium azide, the control vehicle (normal saline) was matched to the same final sodium azide concentration. Throughout the 12-week experimental period, food consumption and body weight were recorded. For each cage, we determined food intake as the difference between the amount of food provided and the remaining food at the end of the week. Body weight was also measured weekly using a digital balance. The experimental procedures were approved by the Institutional Animal Care and Use Committee of The University of North Carolina at Greensboro (IACUC #2024-1212) and carried out in accordance with the ARRIVE 2.0 guidelines.

### 2.2. Animal dissection and tissue collection

At the end of the experimental period, mice were deeply anesthetized using isoflurane, and blood was withdrawn via cardiac puncture using a sterile syringe and needle. Blood was mixed with 48 μL ethylenediaminetetraacetic acid (EDTA) solution, and 120 μL was directly loaded into the VetScan HM5 Hematology Analyzer (Abaxis, Inc., Union City, CA, USA). The remaining blood was centrifuged at 3000 × g for 10 minutes at room temperature to isolate serum, which was stored at -80 °C. Following this, animals were dissected and perfused with PBS, and the liver was harvested and stored at -80 °C. Animal anesthesia, cardiac puncture, and dissection were performed as per the institutional IACUC guidelines.

### 2.3. Heart Dissection, aortic Root Oil Red O staining and Image J analysis

Following animal dissection, we perfused hearts of C57BL/6, LDLr⁻/⁻ and LDLr⁻/⁻+MPLs groups with 1X PBS. For preparation of aortic roots, the heart and proximal aorta were dissected, and the heart was separated from the aorta by a close cut to the base. Using a scalpel, ∼ 70% of the ventricles (from the apex to ∼ 3 mm below the base) were removed, leaving the aortic root containing the upper portion. The aortic portion was embedded in OCT compound in a mold. The aortic root tissue in the OCT compound was mounted on Frozen cryostat chucks, and sectioning was performed through the ventricular tissues and discarded until the aortic sinuses were reached. The aortic sinus was identified by the appearance of all three aortic valve cusps. Following that, serial cryosections of 15 µm were collected (typically 4-5 per slide) until the aortic wall disappeared.

Fresh oil red O (ORO) solution was prepared by mixing 6 mL of saturated ORO stock (0.25-0.5% in isopropyl alcohol) with 4 mL ddH2O, allowing it to stand for 5 mins, and then filtering through filter paper. 10% neutral buffered formalin solution was used to fix slides for 5 min at room temperature, blotted, and then equilibrated in 60% isopropyl alcohol for 5 min at room temperature, and then stained with filtered ORO for 10 min at room temperature. Slides were then treated with 60% isopropyl alcohol for 2 min, rinsed once in dH2O, and counterstained with hematoxylin for 10 seconds. These slides were then rinsed 3 times in dH2O and mounted in warmed glycerol-gelatin. For microscopy, brightfield microscopy at 4x magnification and a 500 µm scale bar was used, keeping all the settings constant across all groups (C.F.A. Culling 1985).

All images were analyzed in Fiji (ImageJ v1.54g) by defining intima. Lumen to internal elastic lamina (IEL) annulus or region was traced and combined with the XOR feature of Fiji, yielding the Intima ROI. The wand tool of the image was used wherever needed, with a tolerance of 10. Color deconvolution isolated the red chromogen, and OTSU was used to determine the most informative threshold channel, producing a binary mask with white indicating ORO-positive cells. We measured % ORO-positive areas with an intima ROI applied to the mask and the limit threshold enabled. A one-way ANOVA followed by Tukey’s HSD at α = 0.05 was applied to all groups.

### 2.4. RNA Extraction, Library Preparation and Sequencing

For RNA Sequencing, RNA was isolated, and an on-column DNase treatment was performed using a Direct-zol RNA miniprep kit (Cat# R2050, Zymo Research). The RNA from 3 mice was randomly pooled to yield 1 biological replicate per group (n=3 per group). Only high-quality RNA was advanced to library preparation following an RNA quantity and quality check using Nanodrop and then Agilent 2100 Bioanalyzer. Poly(A)+ RNA was converted to cDNA following a purification with oligo-(dT) magnetic beads and fragmentation in divalent cation buffer at elevated temperature. Illumina TruSeq Stranded mRNA libraries were prepared, and quality control was performed using an Agilent High-Sensitivity DNA chip (LC Sciences, Houston, TX, USA). Paired-end sequencing was performed using Illumina NovaSeq 6000 using the manufacturer’s guidelines.

### 2.5. Read processing, alignment, assembly, and data analysis

Cutadapt v1.10 was used to remove adapters and poor and low-quality reads (Martin 2011). The quality of the remaining sequences was inspected using FastQC v0.10.1. High-quality and cleaned reads were aligned to Ensembl V112 using HISAT2 v2.0 with default splice-aware parameters (Kim, Paggi et al. 2019). Assemblies for each sample were generated using StringTie v1.3.4 (Pertea, Pertea et al. 2015). To create a unified reference, assembled GTFs were merged using gffcompare, and expression was re-estimated against the merged annotation with StringTie/Ballgown. Following this, gene-level counts were generated from StringTie outputs using prepDE.py to obtain raw gene counts for differential testing, and transcript-level abundance was taken from StringTie/Ballgown tables as FPKM for isoform-level statistics and plotting. The reference resources are Genome: *Mus musculus* Ensembl V112 (GRCm39) DNA FASTA and matching GTF; FTP: ftp://ftp.ensembl.org/pub/release-112/fasta/mus_musculus/dna/.

### 2.6. Differential analysis and data visualization

Differential expression was performed on raw gene counts using DESeq2 with median ratio normalization, and Wald tests (Love, Huber et al. 2014), and Benjamini-Hochberg padj was used for multiple testing with a significance defined at padj < 0.05. A threshold of |log2FC|≥0.58 was used for volcano plot, whereas significance (discovery counts) reported in results was defined by padj (padj < 0.05). For group comparisons, DESeq2 with n=3 pooled replicates per group was used. StringTie/Ballgown FPKM tables with padj < 0.05 were used to define transcript-level (isoform) significance. For any isoform meeting this threshold, we considered the corresponding gene to have ≥ 1 significant transcript. Following analysis, retained genes (22,448) and transcripts (76,694) were visualized with variance-stabilizing transformation (VST) PCA plots, volcano plots (|log₂FC| > 0.58, padj < 0.05), three-way DEG overlap Venn diagrams of significant genes (padj < 0.05), and a heatmap (Z-score-scaled log₂ (FPKM + 1)). For atherosclerosis heatmap, a 502 unique gene pool was made using KEGG mmu05417 (fluid shear stress and atherosclerosis), and mmu05418 (chemical carcinogenesis, reactive oxygen species) and mmu04932 (non-alcoholic fatty liver disease) mapped through org.Mm.eg.db (v3.21.0) with a literature list which covers lipid metabolism, foam cell biology, inflammation and oxidative stress signaling, endothelial function, fibrosis, cell death, cholesterol biosynthesis, and xenobiotic metabolism. Over-representation analysis was performed against GO Biological Process, GO Molecular Function, GO Cellular Component, and KEGG using Benjamini-Hochberg correction (padj < 0.05), and the top 15 terms were displayed. Gene set enrichment analysis was performed on gene data against MSigDB Hallmark and KEGG gene sets (msigdbr v26.1.0, Mus musculus; size filter 15–500 genes).

### 2.7. Alternative Splicing, isoform switch, and prediction of novel transcripts

Replicate-aware alternative splicing (AS) was done using rMATSv4.1.1 (Shen, Park et al. 2014) across five events, i.e., skipped exon (SE), retained intron (RI), mutually exclusive exons (MXE), A5SS, and A3SS, and events were classified as significant at padj < 0.05, |ΔPSI| ≥ 0.2. For sashimi visualization, rmats2sashimiplot was used on SE at padj-significant events. Significant AS events (padj < 0.05, SE, RI, MXE, A5SS, A3SS) were displayed as bar graphs, and sashimi plots of representative SE events were generated using rmats2sashimiplot (padj < 0.05), showing read coverage (RPKM). Within the same gene across different transcripts, we defined an isoform switch as ≥ 1 significantly upregulated and ≥ 1 significantly downregulated transcripts at padj < 0.05. Isoform usage bar plots were generated for the significant transcripts only, with bars indicating mean FPKM per group. To emphasize reciprocal usage, transcripts are ordered from top to bottom. UpSet was created to visualize and quantify overlaps among genes with significant p-values and genes with ≥ 1 significant transcript. **Table S1** provides transcripts-per-gene switch calls and transcript IDs, and **Table S2** highlights genes that are not significant at the gene level but have ≥ 1 significant transcript. **Table S3** contains atherosclerosis-relevant switch candidates. Representative isoform switching events were visualized through bar plots displaying mean FPKM ± SEM.

### 2.8. Comparative transcriptomics analysis with human atherosclerosis

Bulk-RNA Seq data human atherosclerosis data [GSE100927, 104 human atherosclerotic peripheral artery samples (carotid, femoral, infra-popliteal) and non-atherosclerotic controls] were compared to our mouse hepatic transcriptomics data to identify common atherosclerosis signatures. DEGs in human atherosclerosis data were identified using limma (v3.54) at padj < 0.05. Mouse symbols were converted to human orthologs using the NCBI HomoloGene database by retaining only strict 1:1 ortholog pairs (mouse taxonomy ID 10090 and human 9606 joined on shared HID). Directional concordance was determined for each overlapping gene by comparing the log2FC signs across both data sets. It was represented in scatter plots of mouse log2FC (x-axis) versus human log2FC (y-axis), with a linear regression line. GO Biological Process enrichment was performed using all expressed human genes from GSE100927 as the background for the overlapped gene set identified in the LDLr⁻/⁻+MPLs vs C57BL/6 comparison. After converting HGNC gene symbols to Entrez IDs using org.Hs.eg.db and KEGG pathway enrichment were carried out. Multiple testing was performed using the Benjamini–Hochberg correction, with significance defined as padj < 0.05.

### 2.9. Metagenomics Methods

Microbial DNA was extracted from colonic fecal samples using a commercial R1051 isolation kit (Zymo Research, Irvine, CA). The V4 hypervariable region of the 16S rRNA gene was amplified from 50 ng of genomic DNA using 515F/806R primers. Following amplification, amplicons were pooled and sequenced on an Illumina MiSeq platform (LC Sciences, Houston, TX, USA). Bioinformatics processing of 16S rRNA gene data to produce ASV tables: QIIME2 was used to process raw paired-end FASTQ reads (Bolyen, Rideout et al. 2019). Forward and reverse reads were merged, demultiplexed, filtered for quality, chimeras removed, and denoised with DADA2 (Callahan, McMurdie et al. 2016), yielding clean, high-quality amplicon sequence variants (ASVs). Venn diagrams were constructed based on the presence or absence of ASVs in each group to visualize their distribution. All downstream analysis was based on the ASVs abundance table.

Alpha diversity (within-sample diversity) analysis, including Chao1, ACE, observed ASVs (richness), Shannon and Simpson diversity, and Pielou’s evenness and Good’s coverage (sampling completeness), was estimated from the ASV tables. Violin plots were used to visualize Alpha diversity, and group differences were assessed using nonparametric tests, including the Wilcoxon rank-sum test for pairwise comparisons and the Kruskal-Wallis test for multi-group comparisons, as appropriate.

We used taxonomy and phylogeny-based distance metrics to quantify between-sample (Beta) diversity from ASV tables. Distance matrices were computed using Bray-Curtis and Jaccard dissimilarities, unweighted and weighted UniFrac distances, which incorporate phylogenetic information. The analysis includes Principal components analysis (PCA), Principal coordinate analysis (PCoA), non-metric multidimensional scaling (NMDS), and UPGMA hierarchical clustering. The difference in community structure among groups was tested using analysis of similarities (ANOSIM) and permutational multivariate analysis of variance (PERMANOVA/adonis) on distance metrics.

For taxonomic analysis, SILVA (release 138), and NT-16 with a confidence level set to be >0.7 (Quast, Pruesse et al. 2013) was used. Taxonomic assignments were obtained at all levels (Domain, Phylum, Class, Order, Family, Genus, and Species). For the generation of the taxon-level abundance table, ASV counts were collapsed at each taxonomic level. Finally community composition were visualized and summarized using stacked bar plots (showing top 20-30 most abundant taxa), Heatmaps (showing top 30 taxa with hierarchical clustering of taxa), Venn diagram (demonstrating shared and unique taxa), Bubble plot (where size of the bubble shows relative abundance and color denoted taxonomic rank or group), Circos plots (showing the contribution of dominant taxa to each group and distribution of dominant taxon across groups), Sankey plot (showing the relative abundance of top 10 in abundance flora at the phylum and genus level corresponding to different groups (shown on the left) hence presenting the taxa annotation information, corresponding to relationship, proportional shared of the two levels), and Phylogenetic tree (visualizing evolutionary relationships among dominant genera).

### 2.10. Differential abundance and biomarker analysis

We performed differential abundance at the phylum and genus levels by prior removal of low-abundance taxa. Differences between different groups were assessed using an appropriate statistical test, and P values were adjusted for multiple testing. Significant (p < 0.05) taxa were visualized using bar, volcano, and Manhattan plots. Additionally, we also performed indicator species analysis to identify taxa (biomarkers of great significance) that were significantly associated with specific groups. Here, we analyzed the Top 30 most abundant species to identify species that can serve as biomarkers for different groups. We also calculated pairwise Spearman correlation coefficients among the Top 30 genera to investigate relationships among dominant taxa. We visualized them in heatmaps showing the direction of correlation and the size of circles indicating the strength of the correlation. Significance level was applied to all the correlations. Taxa that are favorably or negatively associated with one another can be identified through screening under specific circumstances, which may be essential for revealing their biological importance.

To infer the functional potential of microbes, we used Phylogenetic Investigation of Communities by Reconstruction of Unobserved States (PICRUSt2) (Douglas, Maffei et al. 2020), and functional annotation was performed against the KEGG (Kyoto Encyclopedia of Genes and Genomes) database.

### 2.11. Comparative metagenomics analysis of a human ACVD cohort and mouse nanoplastics exposure

We compared our metagenomics data with the shotgun metagenome-wide association study by Jie et al. (2017), which compared 218 patients with atherosclerotic cardiovascular disease (ACVD) and 187 healthy controls from the human population (Jie, Xia et al. 2017). Technical variation between the two studies was removed from the combined genus-level abundance matrix using MMUPHin (v1.20) (Ma, Shungin et al. 2022), and DESeq2 (v1.46) was used to test for differential abundance between ACVD patients and controls, with padj < 0.05 (Love, Huber et al. 2014). Then, 18 shared genera were identified as nanoplastics-associated ACVD dysbiosis because they were significantly different between LDLr⁻/⁻+MPLs mice and human ACVD patients compared with their respective controls. Each of the 18 genera was manually investigated for its involvement in gut microbiome pathways linked to atherosclerosis. Pathway effects were scored as +1 for pro-atherogenic activity, -1 for protective, and 0 when genus-level evidence was insufficient. To understand the involvement of shared genera in cardiovascular diseases, a comparison of these 18 genera across various cardiovascular diseases was conducted using PubMed and Google Scholar. Evidence was classified as Strong (genus reported by ≥ 2 independent human studies), Good (genus reported by at least one human study), Indirect (family level or mechanistic/animal evidence, and No evidence for the 18 shared genera. All figures were generated in R (v4.5.0) (Team 2025) using ggplot2 (tidyverse v2.0.0) (Wickham, Averick et al. 2019).

### 2.12. Integrative transcriptomics and metagenomics analysis

DEGs (p < 0.05) from our nanoplastics-exposed mice were transformed using log(x+1) and analyzed using WGCNA, unsupervised hierarchical clustering with biweight midcorrelation, and co-expression module detection (Langfelder and Horvath 2012). To measure overall module activity, module eigengenes were computed, and Welch’s two-sample t-test was used to compare the module eigengene values between LDLr⁻/⁻ and LDLr⁻/⁻+MPLs groups. Modules with P < 0.05 were displayed and kept for further analysis. We then used GO Biological and KEGG pathway enrichment analysis to evaluate the biological activities of these significantly altered modules. For each module, enrichment was performed using the Benjamini–Hochberg correction (Benjamini and Hochberg 1995), and modules with fewer than 5 mapped genes were excluded. We finally used an association network to integrate hepatic transcriptomic modules with differentially abundant bacterial genera. Each module eigengene was linked to log(x+1)- transformed midcorrelation, and each genus was assigned to the module with the strongest absolute correlation. For visualization, the top 10 bacterial genera per module (by absolute bicor strength) and the top 10 hub genes per module (ranked by module membership score) were selected. The packages used are WGCNA (Langfelder and Horvath 2008), dynamicTreeCut (Langfelder, Zhang et al. 2008), clusterProfiler (Yu, Wang et al. 2012), org. Mm.eg.db, and ggplot2 (Wickham 2016).

### 2.13. LC-MS Analysis of Mice Liver Extracts

A total of 30 mg of frozen mice liver was added to a pre-chilled 2 mL screw cap tube containing five ceramic beads (2.8 mm). A total of 450 µL cold (−20 °C) extraction solvent (MeOH:ACN:H2O) was added to the tube at a ratio of 2:2:1 and was processed in Omni Bead Ruptor Elite at a speed of 5.5 m/s for 3 cycles, 20 seconds each, with 120 seconds rest on ice between cycles. The solution was then equilibrated on ice for 30 minutes, followed by centrifugation at 16,000 × g, 4 °C for 10 minutes. 250 µL of clear supernatant was transferred to pre-chilled LC-MS tubes. Machine blanks (processed blanks) were processed in the same manner as the samples to subtract peaks due to the extraction solvent, tube plastics, and ceramic beads.

Biological replicates of mouse liver extracts reconstituted in acetonitrile-water were analyzed in a Thermo Fisher Q Exactive Plus mass spectrometer (Thermo Fisher Scientific, Waltham, MA) coupled with a Waters Acquity ultra-performance liquid chromatography (UPLC) system (Waters Corporation, Milford, MA). A 5 µL aliquot of each sample was injected using a Waters ACQUITY UPLC HSS T3 column (100 Å, 1.8 µm, 2.1 × 50 mm) operated at 40 °C. The chromatographic separation was performed using a multi-step gradient elution. The column was initially equilibrated at 1% B and held isocratically from 0.00 to 2.00 min. A shallow linear gradient was then applied from 2.00 to 4.00 min, increasing the organic content from 1% to 2% B, followed by an extended low-organic hold at 2% B until 4.00 min. A linear gradient was subsequently applied from 4.00 to 14.00 min, increasing solvent B from 2% to 85% to elute moderately to strongly retained compounds. This was followed by a column wash phase, in which the organic content was increased to 99% B at 15.00 min and held until 18.00 min to remove strongly retained species. The system was then returned to initial conditions (1% B) at 18.10 min and held until 22.00 min to allow for column re-equilibration before the next injection. UPLC eluate was directed to the mass spectrometer through a heated electrospray ionization (HESI) source. Data were acquired in both positive and negative ionization modes using full-scan MS and data-dependent MS/MS. Full-scan spectra were collected at a resolving power of 35,000 over an m/z range of 125–1500, with the automatic gain control (AGC) target set to 1×10⁶ and a maximum injection time of 100 ms. For data-dependent acquisition, the three most intense precursor ions detected in each full scan were automatically selected for fragmentation. Tandem mass spectra were acquired at a resolution of 17,500 using an isolation window of 1.0 m/z, an AGC target of 1×10⁵, and a maximum injection time of 50 ms. Fragmentation was achieved using stepped normalized collision energies of 25, 35, and 45 eV. Additional DDA settings included a minimum AGC threshold of 8×10³ for MS/MS triggering, an intensity threshold of 1.6×10⁵, dynamic exclusion enabled for 5.0 s, and isotope exclusion activated (Vidar, Baumeister et al. 2023).

### 2.14. Metabolomics Data Peak Picking and Data Filtering

Raw LC–MS data files were converted to the mzML format and subsequently processed using MZmine (version 6.3) for feature detection and peak extraction (Schmid, Heuckeroth et al. 2023). A series of MZmine processing modules was applied to generate a comprehensive feature table containing a unique feature ID, corresponding m/z values, retention times, and peak areas across all samples. A detailed summary of the MZmine modules employed and their associated parameters is provided in Supplementary Information **Table S4.**

### 2.15. Metabolites Annotation using xMSannotator

xMSannotator (v1.3.2) was used to annotate features (Uppal, Walker et al. 2017). The input data were pre-processed by removing zero-intensity and zero-variance features, collapsing duplicate m/z-RT values, removing features missing in more than 50% of samples, and imputing remaining missing values using the half-row minimum. The pipeline assigns a composite confidence score (0–3) to each feature-annotation pair by integrating four sequential evidence layers: adduct and isotope network analysis, weighted gene co-expression network analysis (WGCNA)-based co-expression module assignment, metabolic pathway scoring, and Human Metabolome Database (HMDB) mass matching. **Table S5** contains the post-annotation filtering pipeline and all annotation settings. The 80,622 raw feature-annotation pairs were limited to the major M+H Adduct (8,705 non-redundant features), filtered to a confidence score of ≥ 2 (11,990 pairs), and analyzed for differential abundance. Three pairwise two-sided Welch’s t-tests (LDLr⁻/⁻vs C57BL/6; LDLr⁻/⁻+MPLs vs C57BL/6; LDLr⁻/⁻+MPLs vs LDLr⁻/⁻) and one-way analysis of variance (ANOVA) across all three groups were used to evaluate differential abundance. The Benjamini-Hochberg false discovery rate method was used for multiple testing correction, and effect sizes were represented as log2FC. For ANOVA-significant features, the grouped heatmap displays z-score-normalized group-mean log₂ intensities. Rows are grouped according to Ward’s linkage, and columns are arranged according to experimental condition. Box plots show log2(intensity + 1) for some important metabolites, with each panel annotated with all three pairwise p-values (*** p < 0.001, ** p < 0.01, * p < 0.05, ns).

### 2.16. Data, codes, and software availability

Raw and processed RNA-seq data (with pooling metadata) will be deposited in a public repository (GEO) upon acceptance. The tools used for data analysis includes FastQC v0.10.1 (quality control); Cutadapt v1.10 (adapter/quality trimming); HISAT2 v2.0 (spliced alignment); StringTie v1.3.4 (assembly/quantification); DESeq2 (differential expression); rMATS v4.1.1 (alternative splicing). Plotting and composites were generated in R (v4.5.0) and Python (3.14) using standard packages (DESeq2/Ballgown/ggplot2; pandas/numpy/matplotlib).

## Results

### Nanoplastics exposure promotes atherosclerotic lipid accumulation in LDLr⁻/⁻ mice

Representative aortic root cross sections stained with Oil Red O (**Fig.1 A1-A3)** showed an increase in intimal neutral lipid across groups, with little staining in the C57BL/6 group (**Fig.1 A1)**, more extensive deposits in LDLr⁻/⁻ (**Fig.1 A2)**, and the greatest, confluent staining in LDLr⁻/⁻+MPLs group (**Fig.1 A3)**. The intima (lumen-to-IEL annulus) served as the analysis region; images were acquired at 4× with a 500-µm scale.

**Fig. 1.**
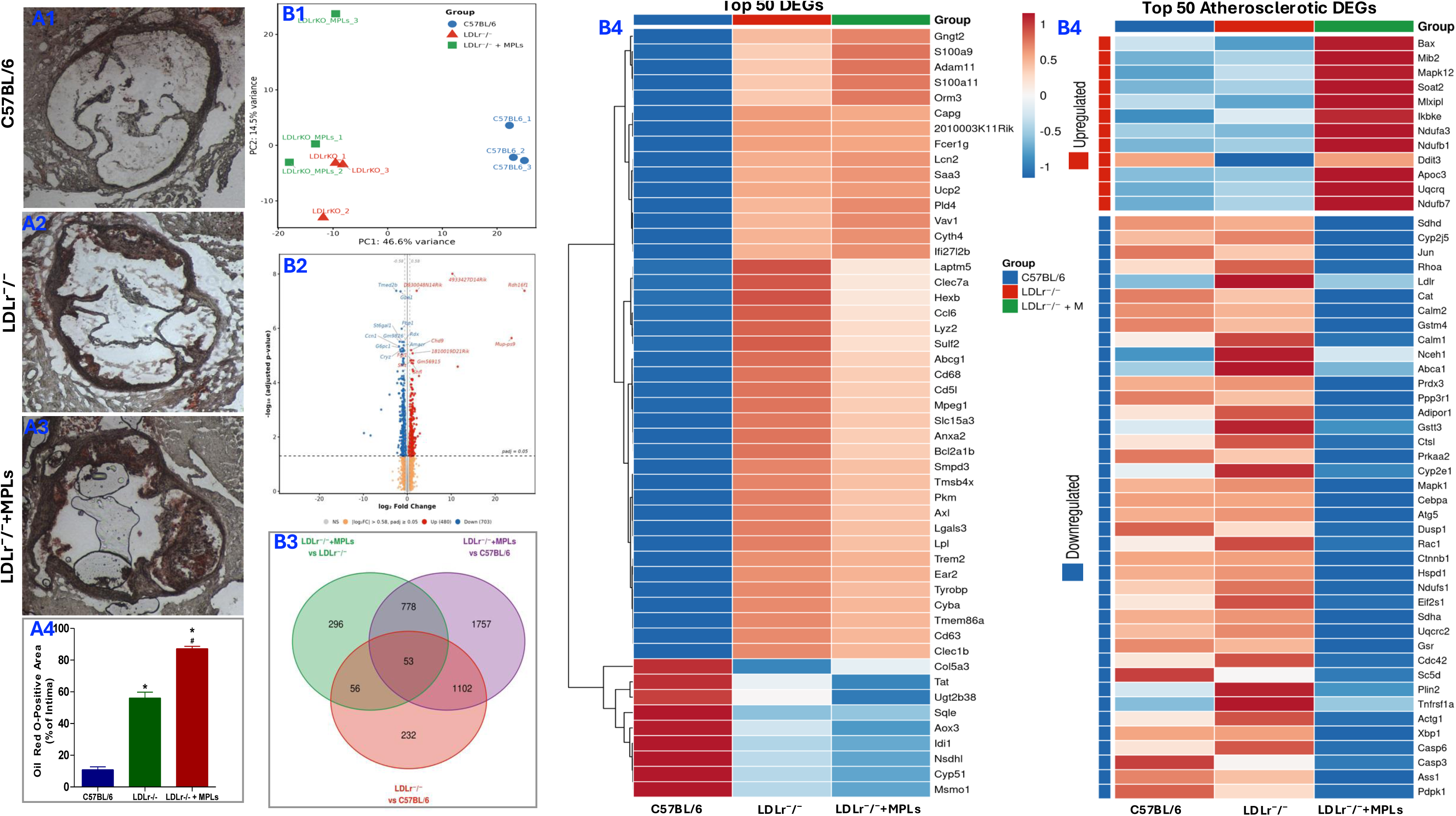
Panel A . Oil Red O staining and quantification of intimal lipid in aortic cross-sections. (A1-A4) Representative sections stained with Oil Red O and counterstained with hematoxylin from C57BL/6 (A1), LDLr⁻/⁻ control (A2), and LDLr⁻/⁻+MPLs (A3). Scale bar: 500 µm. (A4) Quantification of Oil Red O–positive area, expressed as the percentage of intimal area, calculated per animal across all sections. Bars represent mean ± SD with individual animals overlaid (n = 6). One-way ANOVA followed by Tukey’s HSD for post-hoc comparisons, all pairwise group differences are significant, * *p* < 0.05 compared to the C57BL/6 group and # *p* < 0.05 compared to the LDLr⁻/⁻ group. **Panel B. Microplastic exposure induces a predominantly suppressive hepatic transcriptional program in LDLr⁻/⁻ mice.** (B1) Principal component analysis (PCA) of gene expression profiles from liver samples of C57BL/6 mouse (blue, n = 3), LDLr⁻/⁻ mouse (red, n = 3), and LDLr -/- mouse + MPLs (green, n = 3). (B2) Volcano plot of differential gene expression in LDLr⁻/⁻+MPLs versus LDLr⁻/⁻ controls (padj < 0.05). (B3) Three-way Venn diagram of differentially expressed genes across all pairwise contrasts (B4) Heatmap of the top 50 differentially expressed genes (DEGs) (group means). (B5) Heatmap of significantly dysregulated atherosclerosis-related genes (padj < 0.05). Data were generated from liver tissue of LDLr⁻/⁻ mice exposed to polystyrene Nanoplastics (200 mg/kg body weight, 12 weeks; n = 9) or LDLr−/− controls (n = 9), with C57BL/6 mice as reference (n = 9). Each biological replicate represents pooled samples from three mice.

Quantitative analysis of Oil Red O-positive area (% of intima) confirmed these differences. Mean ± SEM values were 10.89 ± 3.58% for C57BL/6, 56.13 ± 8.90% for LDLr⁻/⁻, and 87.17 ± 3.84% for LDLr⁻/⁻+MPLs (n = 6) (**Fig.1 A4)**. One-way ANOVA followed by Tukey’s HSD demonstrated that all pairwise comparisons were significant (C57BL/6 vs LDLr⁻/⁻, P < 0.001; C57BL/6 vs LDLr⁻/⁻+MPLs, P < 0.001; LDLr⁻/⁻ vs LDLr⁻/⁻+MPLs, P < 0.001). These results indicate that LDLr deficiency markedly increases intimal lipid relative to normal controls and that exposure further increased lipid accumulation beyond the LDLr⁻/⁻ baseline.

### Nanoplastics reprogram the transcriptome profile of LDLr⁻/⁻ mice liver

The liver transcriptome of LDLr⁻/⁻ mice exposed to polystyrene nanoplastics (LDLr⁻/⁻+MPLs) and LDLr-control (LDLr⁻/⁻), and normal-control (C57BL/6) was profiled. PCA showed a tight within-group clustering and clear separation between the three groups along PC1 (46.6% variance) and PC2 (14.5% variance) (**Fig. 1 B 1)**. 1,183 differentially expressed genes (DEGs, padj < 0.05) with 480 upregulated and 703 downregulated between LDLr⁻/⁻+MPLs vs LDLr⁻/⁻ (**Fig.1 B2)**, demonstrating a transcriptional response to nanoplastics exposure. 296 DEGs are specifically dysregulated by nanoplastics exposure (absent from both the disease-baseline and combination contrasts), following comparison of all three pairwise contrasts (LDLr⁻/⁻+MPLs vs. LDLr⁻/⁻, LDLr⁻/⁻+MPLs vs. C57BL/6, and LDLr⁻/⁻ vs. C57BL/6) (**Fig.1 B3**; directional Venns for upregulated and downregulated subsets in **Fig. S1 C–D**). A heatmap of the top 50 DEGs from all contrasts shows a clear separation and distinct expression pattern for all DEGs (**Fig.1 B4, Fig. S1 E).** An analysis of a curated atherosclerosis gene pool identified 52 significantly dysregulated genes related to atherosclerosis, with 40 (76.9%) exhibiting downregulation and 12 (23.1%) upregulation, resulting in a suppression-to-activation ratio of 3.3:1. The downregulated genes included mediators of cholesterol biosynthesis (*Cyp51, Sqle, Idi1*), reverse cholesterol transport (*Abca1, Ldlr*), mitochondrial function (*Sdhd, Prdx3*), and cardioprotective lipid-mediator synthesis (*Cyp2j5*). The upregulated genes included the ER-stress marker *Ddit3*, the pro-apoptotic factor *Bax*, and mitochondrial complex subunits (*Ndufa3, Ndufb7, Uqcrq*) (**Fig. 1B4, Fig. S1F**). Volcano plots for the remaining two comparisons are provided in **Fig. S1A** (LDLr⁻/⁻+MPLs vs C57BL/6) and **Fig. S1B** (LDLr⁻/⁻ vs C57BL/6), and transcript-level volcano plots for all three contrasts in **Fig. S3 A–C**. At the transcript level, differential expression testing across 76,694 isoforms identified 2,297 differentially expressed transcripts in the same contrast (863 upregulated and 1,434 downregulated; padj < 0.05; **Fig. S3B**), reinforcing the predominantly downregulatory signal observed at the gene level.

### Nanoplastics reshape the hepatic transcriptome at the transcript level and induce widespread alternative splicing

We first compared differential expression of genes versus transcript resolution of features that were significant only at the transcript level, only at the gene level, or at both (**Fig. 2A**). Out of all the significant loci (padj < 0.05), 294 genes were significant both at the gene and transcript levels, 129 at the gene level only, and 1927 at the transcript level only, giving a total of 2221 genes with ≥1 significant transcript (**Fig. 2A**). These counts reflect differential transcript expression and indicate that a substantial fraction of the response is detectable only at transcript resolution, they do not, by themselves, distinguish changes in transcript abundance from changes in splicing. To directly assess splicing, we next applied rMATS. Consistent with this, PCA showed clear separation of LDLr⁻/⁻+MPLs samples from both C57BL/6 and LDLr⁻/⁻ groups along PC1 (**Fig. 2B**), confirming a distinct nanoplastics-specific profile at the transcript level.

**Fig. 2.**
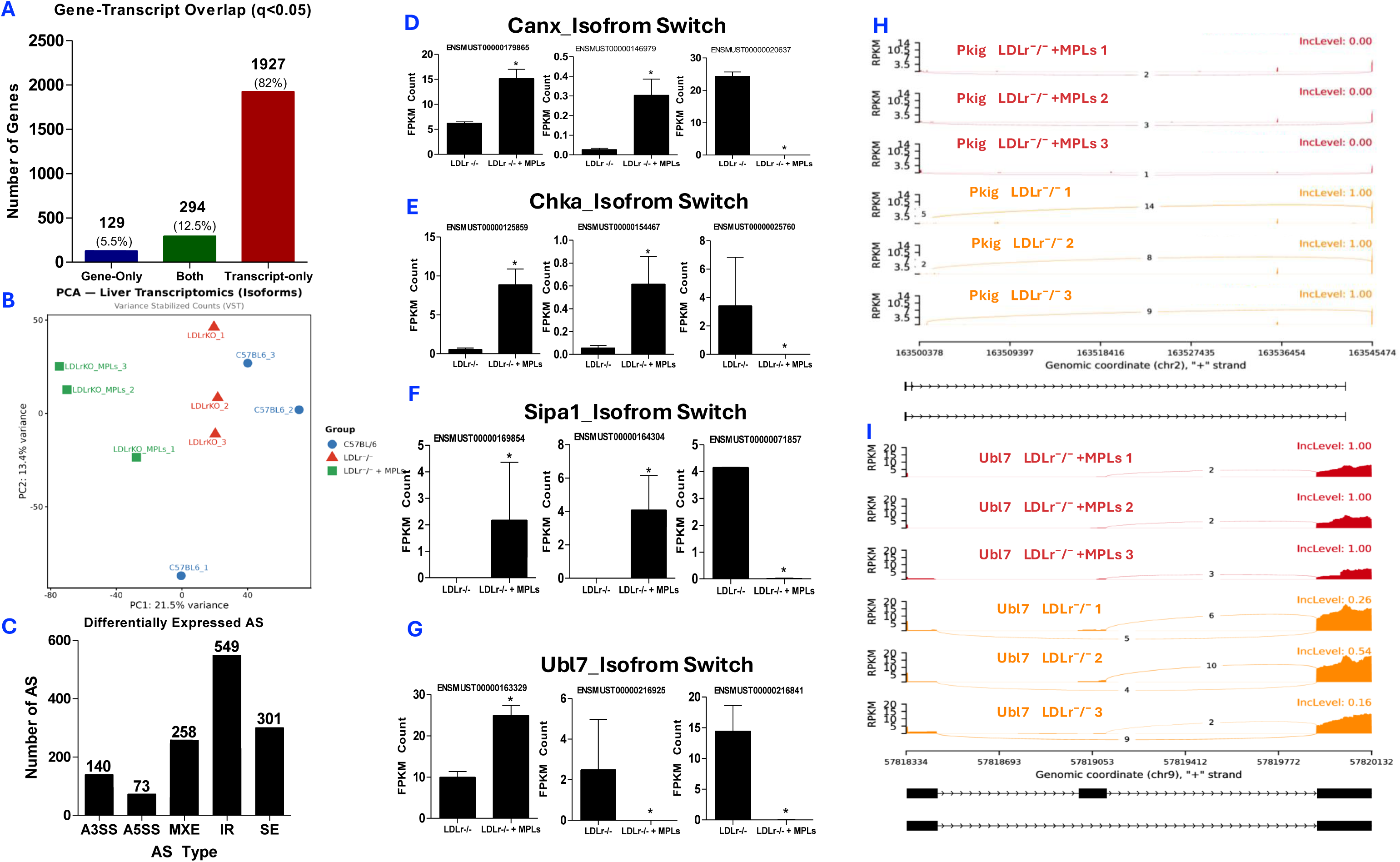
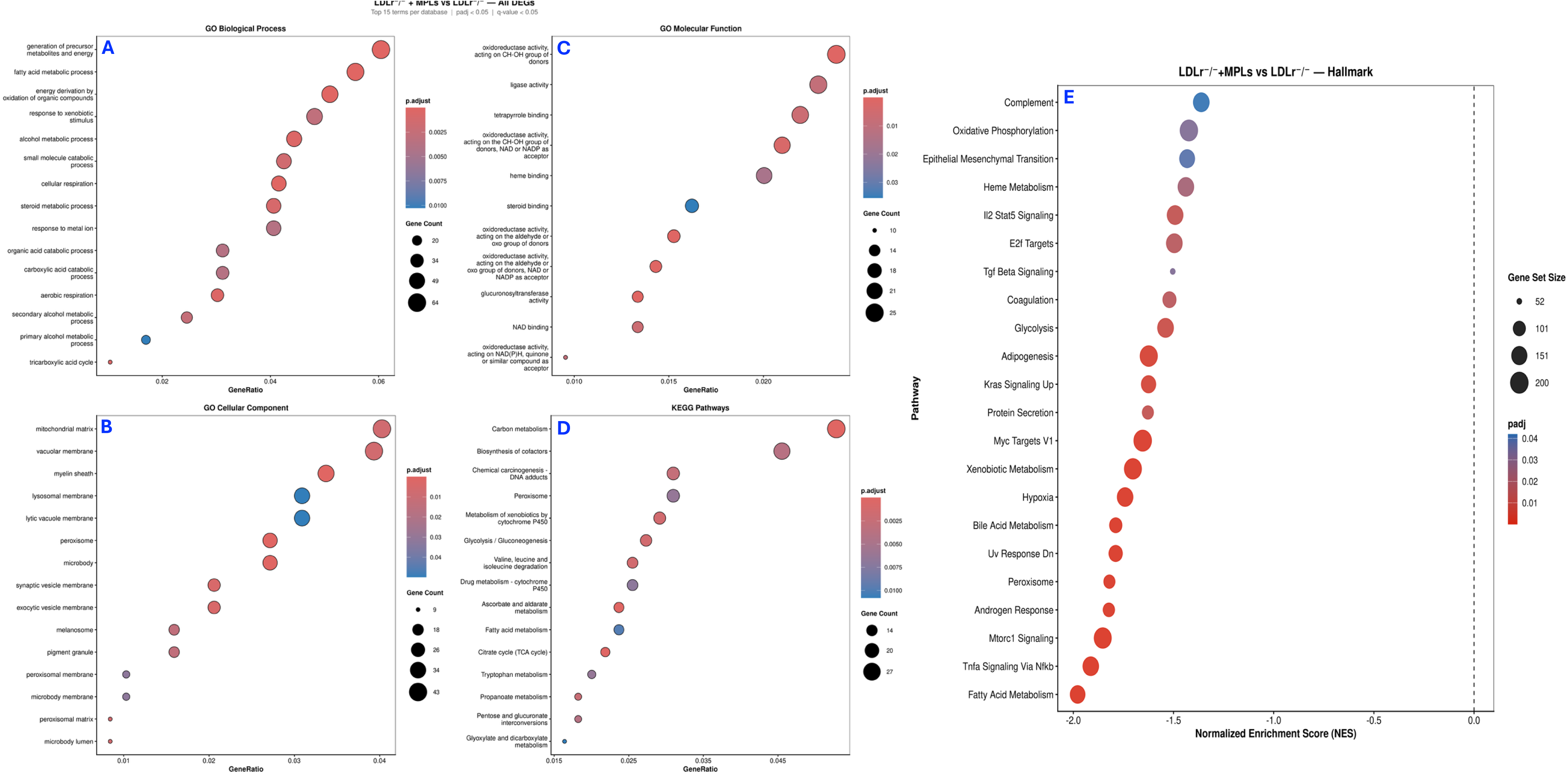
Microplastic exposure induces widespread isoform-level reprogramming and alternative splicing in the atherosclerotic liver. (A) Bar chart showing the overlap between gene-level and transcript-level differential expression (q < 0.05) in the LDLr⁻/⁻+MPLs versus LDLr⁻/⁻ control. (B) Principal component analysis (PCA) of isoform-level expression across C57BL/6 mouse, LDLr⁻/⁻, and LDLr⁻/⁻+MPLs groups. (C) Distribution of significantly altered alternative splicing events identified by by rMATS (q < 0.05), with retained intron (RI) events being the most abundant, followed by skipped exon (SE), mutually exclusive exon (MXE), and alternative splice site events (A3SS, A5SS), indicating widespread disruption of splicing regulation. ((D–G) Representative isoform switching events across key pathways, showing differential transcript usage between LDLr⁻/⁻ control and LDLr⁻/⁻+MPLs. Bar plots display mean expression (FPKM ± SEM) of individual transcript isoforms. (H–I) Sashimi plots provide read-level validation of alternative splicing events for selected genes in LDLr⁻/⁻+MPLs (red) vs LDLr⁻/⁻ control. All biological replicates (n = 3 per group) represents pooled samples from three mice.

To directly quantify splicing rather than transcript abundance, 1321 significant alternative splicing (AS) events were identified between LDLr⁻/⁻+MPLs and LDLr⁻/⁻ using rMATS at padj < 0.05,|ΔPSI| ≥ 0.2. Retained intron (RI) were most frequent (41.56%, 549/1321), followed by skipped exons (SE, 22.8% 301/1321), mutually exclusive exons (MXE, 19.5%, 258/1321) and alternative 3’ splice sites (A3SS, 10.6%, 140/1321) and alternative 5’ splice sites (A5SS, 5.5%, 73/1321) (**Fig. 2C**).

Nanoplastics induce changes in genes at sentinel loci that influence atherosclerosis. For instance, *Agpat3* (1-acylglycerol-3-phosphate O-acyltransferase 3) displayed clear exon inclusion in LDLr⁻/⁻+MPLs (ΔPSI = +0.32, padj = 5.80*10^-12^, **(Fig. S3 G)**, *Dpp9* (dipeptidyl peptidase 9) shows inclusion in LDLr⁻/⁻+MPLs (ΔPSI = +0.189, padj = 2.76*10^-8^, **Fig.S3 E)**, *Ubl7* (ubiquitin-like 7) showed near complete exon inclusion in LDLr⁻/⁻+MPLs vs control (ΔPSI = +0.682, padj = 1.90*10^-8^, **Fig. 2G**, with sashimi validation in **Fig. 2H–I**), *Eif4a2* (eukaryotic translation initiation factor 4A2) showed inclusion in LDLr⁻/⁻+MPLs vs control (ΔPSI = +0.189, padj = 2.76*10^-8^, **Fig. S3 F)**.

Isoform switching was defined as ≥1 up- and ≥ 1 down-regulated transcripts from the same gene (padj < 0.05). A total of 153 genes showed opposite-directional significant transcripts at the defined criteria (**Table S1).** Isoform switch bar plots demonstrate cardiovascular-related diseases, especially atherogenic properties (**Fig. 2D–G; Fig. S3 A–D)**. Representative genes with significant isoforms having roles in diseases or cellular processes disruption includes *Canx* (2↑/1↓) which may have a role in Proteostasis/ER stress because as an ER chaperone, it influences folding and receptor trafficking shifts (**Fig. 2D**), *Sipa1* (2↑/1↓) shows Rap1 dependent adhesion and barrier control and hence may relate to endothelium relevant junctional signaling (**Fig. 2F**), *Cbs* (1↑/3↓) influences H2S/NO linked carbon metabolism **(Fig. S3 D)**, *Chka* (**Fig. 2E**) and *Acbd5* **(Fig. S3 A)** (each with 2↑/1↓) play a role in phospholipid synthesis and in peroxisomal long chain fatty acid transport rewiring hence may play a role in lipid handling, *Men1* (1↑/2↓) **(Fig. S3 B)** and Anapc5 (1↑/3↓) **(Fig. S3 C)** provide support to sustained cellular activation and hence may relate to transcription and ubiquitin cell cycle.

To ensure we were seeing splicing changes rather than just the whole gene going up or down, Isoform switch bar plots were paired with Sashimi plots (**Fig. 2H–I; Fig. S3 E–G).** For example, for *Ubl7*, as shown in the bar plots, one transcript increased and two decreased (a true switch). The splice junction read flipped in the same way, meaning that the sashimi plot shows the expected change in junction usage: more inclusion arcs in LDLr⁻/⁻+MPLs and fewer in controls, which matches the switch (**Fig. 2H–I**). Other representative genes with a plausible role in atherosclerosis, such as *Eif4a2* **(Fig. S3 F)**, *Agpat3* **(Fig. S3 G)**, and *Dpp9* **(Fig. S3 E),** show the same pattern: sashimi plot junctions mirror dominant transcript changes. These demonstrate and support post-transcriptional regulation as a prominent componenet of the hepatic response.

### Nanoplastics affect hepatic metabolic and atherosclerosis-relevant pathways

To investigate hepatic transcriptional pathway changes, enrichment analysis was performed on 1183 DEGs identified between LDLr⁻/⁻+MPLs and LDLr⁻/⁻ group. GO and KEGG analyses revealed significant enrichment terms related to precursor metabolite and energy generation, fatty acid metabolism, oxidative metabolism, xenobiotic response, cellular and aerobic respiration, steroid metabolism, mitochondrial matrix, peroxisome, oxidoreductase activity, heme and tetrapyrrole binding, carbon metabolism, glycolysis/gluconeogenesis, citrate cycle, and cytochrome P450-mediated xenobiotic and drug metabolism (**Fig. 3A–D**). Hallmark GSEA identified 22 gene sets significantly enriched with negative normalized enrichment scores (**Fig. 3E).** These results suggest that nanoplastics exposure disrupts hepatic lipid handling, inflammatory signaling, and xenobiotic-processing pathways, thereby highlighting impairment of metabolic and atherosclerosis-relevant transcriptional programs in LDLr⁻/⁻ mice.

### Microplastic-exposed atherosclerotic liver transcriptome converges with human atherosclerosis

To investigate the role of nanoplastics-induced transcriptional reprogramming in atherosclerosis, mouse DEGs were compared with human atherosclerosis DEGs (data set: GSE100927). A total of 183 genes were shared with the ACVD. The combined ACVD and nanoplastics transcriptomics profile significantly mimics human atherosclerotic gene expression, as shown by the 183 genes shared with the ACVD transcriptomics profile with a highly significant overlap (one-sided Fisher’s exact test, p < 0.001) (**Fig 4A**). Directional concordance analysis of overlapping genes revealed strong cross-species (mice vs. humans) consistency, with the majority of genes concordantly upregulated in both species, including key macrophage and inflammatory markers such as *Trem2, Cd68, Tyrobp, Lpl, Cyth4, Pla2g7, Vav1*, and *Capg* (**Fig. 4 B)**. A few discordant genes (such as *Pi16, Scara5, Des, Hp, Wnt11* up in mouse but down in human, and *Serpina1e, Fbp1, Cd163* down in mouse but up in human) suggest species-specific differences. GO Biological enrichment analysis of the shared 183 genes showed immune activation and inflammatory signaling terms (positive regulation of cytokine production, leukocyte cell–cell adhesion, immune response–regulating signaling pathway, regulation of lymphocyte activation, and leukocyte migration) (**Fig. 4C**). KEGG pathway enrichment identified significant pathways such as Phagosome and Cytokine–cytokine receptor interaction (**Fig. 4D**). These enrichment analyses provide directed pathway-level evidence that transcriptional overlap between nanoplastics treated mouse liver and human atherosclerosis is biologically relevant vascular biology. These results reveal that nanoplastics exposure amplifies and accelerates atherosclerosis-relevant transcriptional changes.

**Fig. 4.**
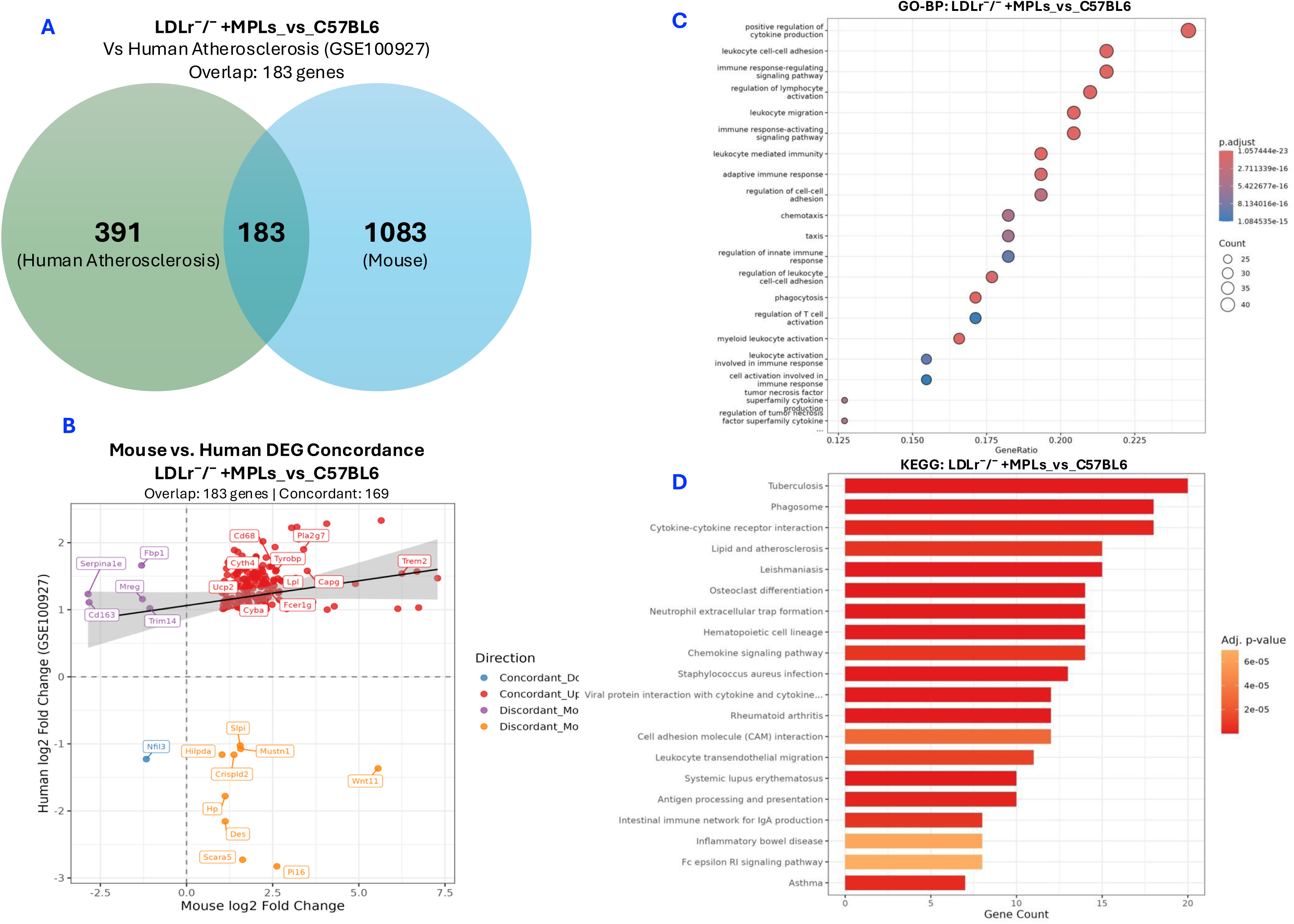
Cross-species concordance reveals shared inflammatory and atherogenic profile between microplastic treated mice and human atherosclerosis. (A) Venn diagram showing the overlap between differentially expressed genes (DEGs) from LDLr⁻/⁻+MPLs versus C57BL/6 mouse mice (converted to human orthologs) and human atherosclerosis DEGs from GSE100927. (B) Concordance scatter plot of overlapping genes comparing mouse (x-axis) and human (y-axis) log₂ fold changes. Representative genes are labeled, and the regression line indicates a positive correlation across species. (C) Gene Ontology Biological Process (GO-BP) enrichment analysis of overlapping genes. (D) KEGG pathway enrichment analysis of overlapping genes. Enrichment analyses were performed with Benjamini—Hochberg correction (FDR < 0.05).

### ASV profiling and Alpha and Beta diversity profiles in response to Nanoplastics exposure

To quantify the exact number of unique and shared microbial sequences across different groups before downstream analysis, we used a Venn Diagram to analyze ASV profiling data. The Venn diagram showed 278 common genera among all groups, hence representing the core microbiome profile. LDLr⁻/⁻+MPLs group contained 999 unique ASVs, LDLr⁻/⁻ group contained 1242 unique ASVs, whereas C57BL/6 group contained 1200 unique ASVs, highlighting distinct microbial profiles in response to nanoplastics exposure, LDLr deficiency, diet, and host background (**Fig. 5A**). Principal component analysis showed separation between groups PC1=11.68% and PC2=9.81%) (**Fig. 5B**).

**Fig. 5.**
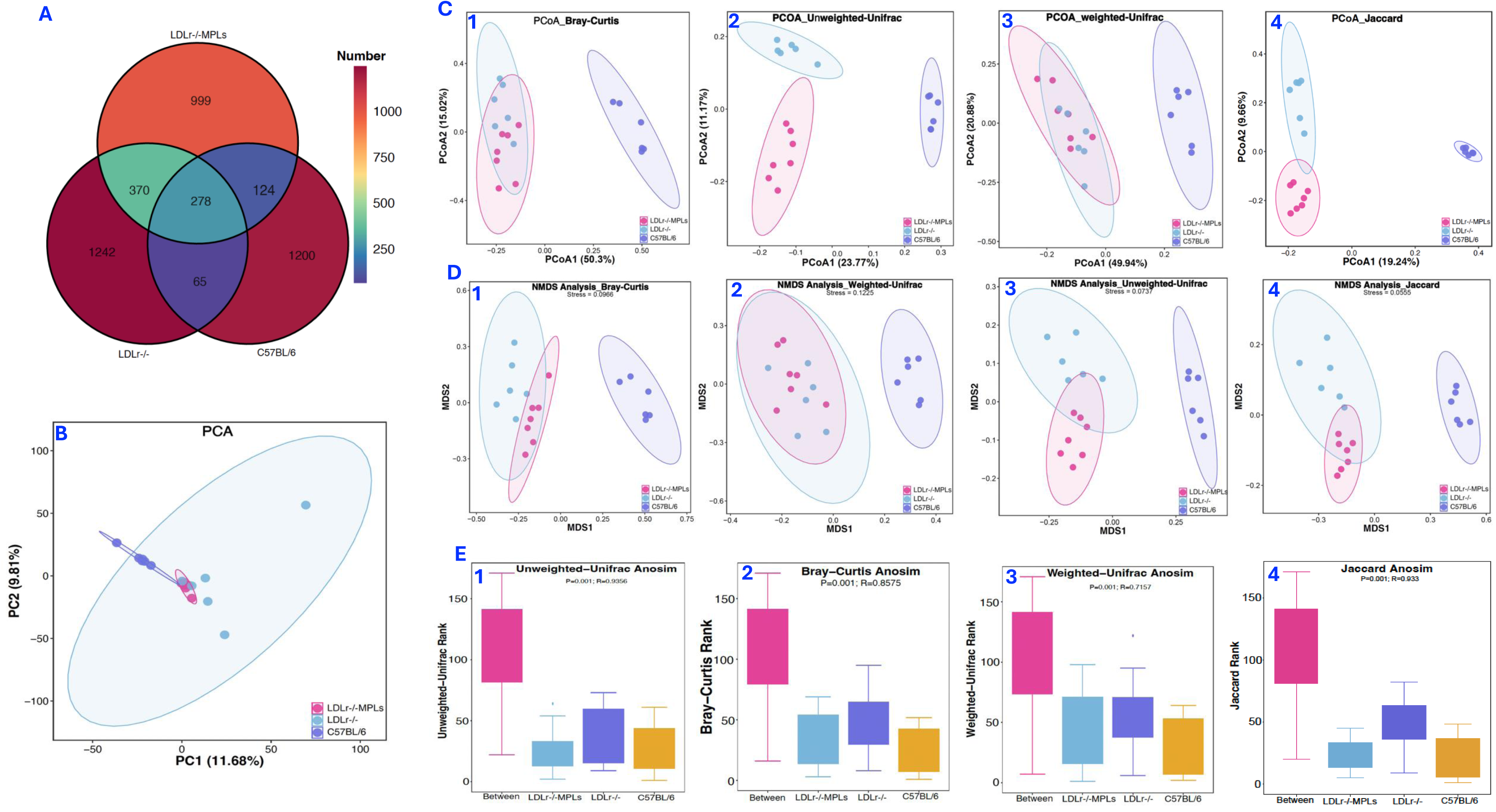
Microplastic exposure induces alterations in the gut microbial community. (A) Venn diagram showing changes in amplicon sequence variant (ASV) profiles across all groups. A total of 278 ASVs were shared among all groups, whereas 999 ASVs were unique to the LDLr⁻/⁻ mouse + MPLs group. (B) β-diversity analysis by principal coordinates analysis (PCoA) demonstrating clear separation among all experimental groups. **(C-E) Microplastic exposure reshapes gut microbial beta diversity in LDLr⁻/⁻ mice.** (A) Principal coordinates analysis (PCoA) based on four distance metrics; Bray–Curtis (PCoA1 = 50.3%, PCoA2 = 15.02%), weighted UniFrac (PCoA1 = 49.9%, PCoA2 = 20.88%), unweighted UniFrac (PCoA1 = 23.77%, PCoA2 = 11.7%), and Jaccard (PCoA1 = 19.24%, PCoA2 = 9.66%), revealed clear separation and distinct clustering among LDLr⁻/⁻ mouse + MPLs, LDLr⁻/⁻, and C57BL/6 mouse groups. (D) Non-metric multidimensional scaling (NMDS) using the same distance metrics further confirmed group-wise separation, with low stress values indicating good ordination fit (Bray–Curtis = 0.0966, weighted UniFrac = 0.1225, unweighted UniFrac = 0.0737, Jaccard = 0.0555). (E) Analysis of similarities (ANOSIM) demonstrated significant between-group differences across all metrics (p = 0.001), with high R values indicating strong dissimilarity among groups.

To evaluate microbial richness and evenness across groups, Alpha diversity analysis was performed. Alpha-diversity indices (observed species, Chao1, ACE, Shannon, Simpson, Pielou’s evenness, and Good’s coverage) did not differ significantly between the LDLr⁻/⁻+MPLs and diet-and genotype-matched LDLr⁻/⁻ groups for any metric (padj > 0.05; **Table S6**), indicating that nanoplastic exposure did not alter overall within-sample richness or evenness. Significant reductions in Simpson diversity and Pielou’s evenness were observed only in the LDLr⁻/⁻+NPLs versus C57BL/6 comparison (padj = 0.04 for both); however, as these groups differ in genotype, diet, and nanoplastic exposure, this difference cannot be attributed specifically to nanoplastics. Notably, the genotype- and diet-only contrast (LDLr⁻/⁻ versus C57BL/6) was non-significant for all indices (padj > 0.05), indicating that the observed evenness and Simpson differences do not arise from genotype or diet alone.

We used multiple distance metrics (Bray-Curtis, Jaccard, unweighted UniFrac, and weighted UniFrac) to perform Beta Diversity analysis for measuring variability between groups. Principal coordinate analysis (PCoA) on Jaccard (PCoA1= 19.24%, PCoA2= 9.66%), weighted UniFrac (PCoA1 = 49.9 %, PCoA2 = 20.9 %), unweighted UniFrac (PCoA1 = 23.77 %, PCoA2 = 11.7 %), and Bray Curtis (PCoA1 = 50.3 %, PCoA2 = 15.02 %), displayed clear separation and distinct clustering pattern among all three groups (**Fig. 5C**). Non-Metric multidimensional scaling (NMDS) further confirmed the separation between groups based on Jaccard (Stress = 0.05), weighted UniFrac (0.1225), unweighted UniFrac (0.0737), and Bray Curtis (0.0966) displayed clear separation and distinct clustering pattern among all the three groups (**Fig. 5D**). The PERMANOVA (adonis) tests further confirmed the statistically significant differences in microbial community composition among all the three Groups (**Fig. 5E**). For Bray Curtis (F= 12.15, R^2^ = 0.603, p = 0.001), and weighted UniFrac (F = 10.98, R^2^ = 0.578, p = 0.001) we observed the strongest effect **(Table S7, adonis combined table)**. Together, these analyses demonstrate that nanoplastics exposure significantly altered the gut microbiome in the LDLr⁻/⁻+MPLs group.

### Taxonomic and differential abundance analyses reveal the role of nanoplastics exposure in atherosclerosis through Gut microbiome dysbiosis

Taxonomic community structure at both the phylum and genus levels was observed across all groups. At the genus level, z-Score heatmap clustering demonstrated enrichment of *Romboutsia*, *Faecalibaculum*, *Oscillibacter*, *Coprococcus*, *Colidextribacter*, *Incertae_Sedis*, *Intestinimonas,* and depletion of *Anaerotignum, Clostridium_sensu_stricto_1, Eubacterium, Erysipelatoclostridium, Eubacterium _nodatum_Group, Eisenbergiella, GCA_900066575, Acetatifactor, Dubosiella, Turicibacter, Roseburia*, and *Bifidobacterium* in the LDLr⁻/⁻+MPLs group relative to LDLr⁻/⁻ (**Fig. 6A**). Venn Diagram demonstrated 97 distinct genera in the LDLr⁻/⁻ group (**Fig. 6B).** Differential abundance was also calculated at the genus level (enriched or depleted, padj < 0.05) in the LDLr⁻/⁻+MPLs group, including the depleted genera such as *Bacteroides, Barnesiellaceae, Christensenellaceae, Marvinbryantia, Eubacterium, Lachnoclostridium, and Flavonifractor* (Fig. 6C-I). The enriched genera include *Streptococcus, Romboutsia*, and *Faecalibaculum* (Fig 6J-L).

**Fig. 6.**
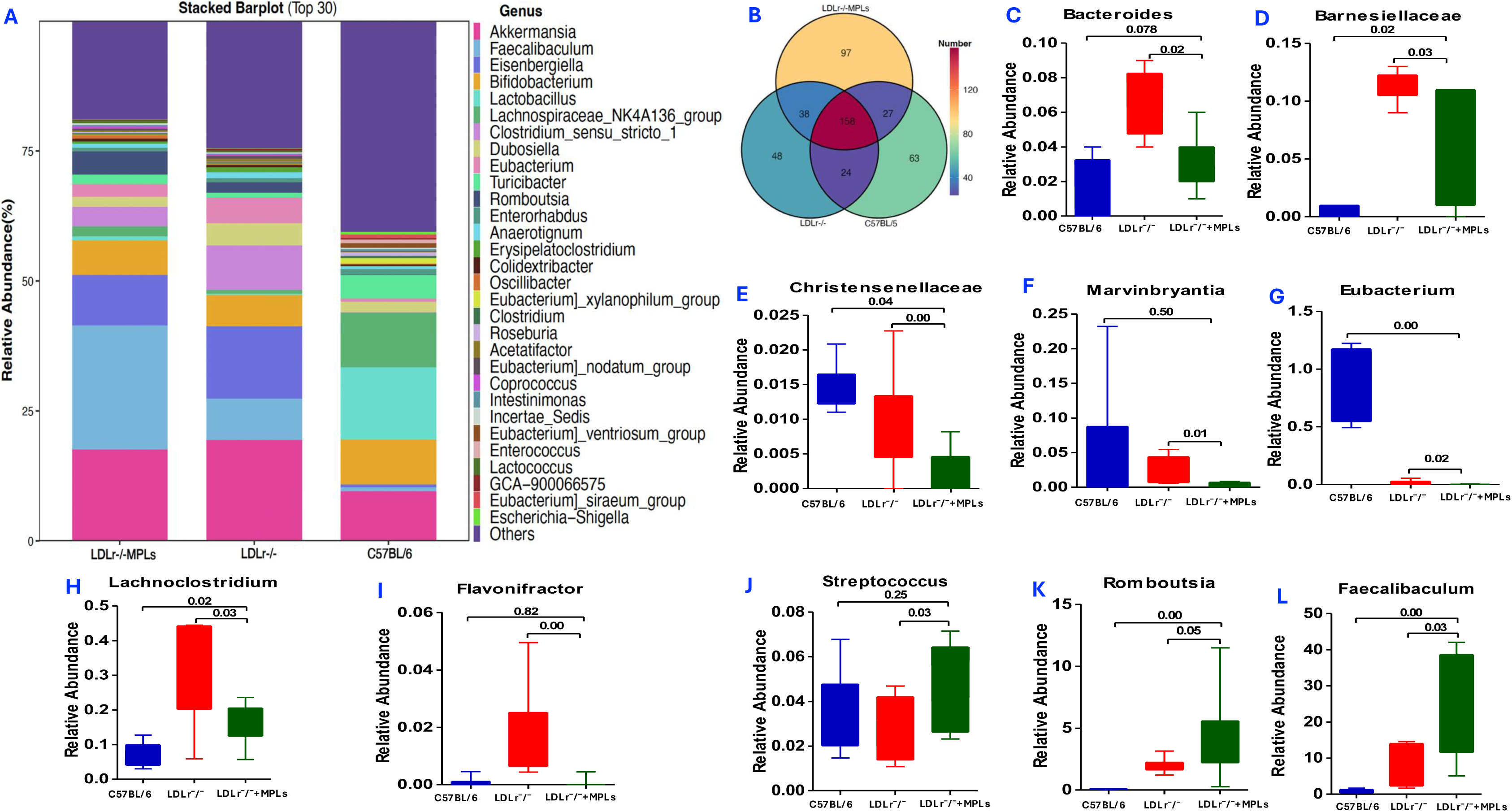
Genus level Taxonomic community shifts and differential microbial abundance in response to microplastic exposure. (A) Heatmap showing normalized relative abundance z-scores of the most dominant bacterial genera across all three groups. Distinct clustering patterns highlight microplastic-induced changes in taxonomic composition. (B) Venn diagram of bacterial genera in different groups (C-L) Genus-level differential abundance analysis of key taxa, including (C) *Bacteroides, (D) Barnesiellaceae, (E) Christensenellaceae, (F) Marvinbryantia, (G) Eubacterium, (H) Lachnoclostridium, (I) Flavonifractor, (J) Streptococcus*, (K) *Romboutsia, and (L) Faecalibaculum.* Significant differences between LDLr⁻/⁻+MPLs and LDLr⁻/⁻ groups demonstrate marked taxonomic remodeling in response to microplastic treatment.

At the genus level, Indicator species analysis revealed distinct group-specific microbiome patterns **(Fig. S4A)**. *Faecalibaculum*, *Romboutsia*, and *Oscillibacter* demonstrated the highest indicator values, followed by *Coprococcus*, *Lactococcus*, *Colidextribacter*, and others **(Fig. S4A)**. *Clostridium_sensu_stricto_1* and *Erysipelatoclostridium* have strong indicator values in the LDLr⁻/⁻ group. In contrast, *Dubosiella*, *Lactobacillus*, *Enterococcus, Turicibacter,* and *Erysipelatoclostridium* showed strong indicator values in the C57BL/6 group **(Fig. S4A)**.

For the phylum-level taxonomy, z-score heatmap clustering showed enrichment of Cyanobacteria, Planctomycota, Firmicutes, Gemmatimonadota, WPS-2, Armatimonadota, Patescibacteria, Deinococcota, Campylobacterota, and depletion of common commensal phyla including Verrucomicrobiota, Proteobacteria, Fusobacteriota, Desulfobacterota, Acidobacteriota, Chloroflexi, and Myxococcota in LDLr⁻/⁻+MPLs groups as compared to the LDLr⁻/⁻ **(Fig. S5A)**. The Venn diagram also showed 2 Phyla that are exclusively present in the LDLr⁻/⁻+MPLs group **(Fig. S5B)**. Differential abundance between groups showed an apparent increase in low-abundance envriomental phyla (Planctomycota and Cyanobacteria) in LDLr⁻/⁻+MPLs as compared to the LDLr⁻/⁻ group **(Fig. S5E-F),** but were not further interpreted.

A Sankey plot summarized the flow of microbial abundance from groups to the phylum and genus levels, demonstrating that the Faecalibaculum genus within the Firmicutes phylum was abundant in LDLr⁻/⁻+MPLs compared to the LDLr⁻/⁻ group. In contrast, the Eisenbergiella genus of the Firmicutes was abundant in the LDLr⁻/⁻ group as compared to the LDLr⁻/⁻+MPLs group. Lactobacillus of the Firmicutes phylum was abundant in the C57BL/6 group as compared to the other two **(Fig. S4B)**.

Manhattan plot clarified the differential abundance of genera between groups in response to nanoplastics exposure. Numerous genera in the phylum Firmicutes, Proteobacteria, Cyanobacteriota, and Actinobacteriota exceeded the significance threshold (−log₁₀P ≥ 2) in the LDLr⁻/⁻+MPLs group relative to LDL-/-. The depletion and enrichment of genera across the phylum are shown in **Fig. S4C.**

### Microbial Co-occurrence Network Reveals Coordinated Dysbiosis in Nanoplastics-Exposed Mice

A correlation matrix analysis of the dominant genera revealed a significant restructuring of microbial co-occurrence networks in LDLr⁻/⁻+MPLs mice, which aligns with the microplastic-induced dysbiosis observed. The plot exhibited distinct clusters of strongly positively correlated taxa, predominantly within the Firmicutes phylum, suggesting coordinated shifts among genera involved in carbohydrate fermentation and short-chain fatty acid (SCFA) metabolism. Conversely, several inflammation-associated genera exhibited strong negative correlations with health-associated commensals, indicating competitive exclusion under nanoplastics exposure. Notably, genera such as Lactobacillus, Akkermansia, and Coprococcus emerged as network hubs with numerous high-magnitude correlations, signifying their central role in shaping community responses. Nanoplastics exposure disrupted these interactions, weakening positive correlations among SCFA-producing genera while strengthening antagonistic relationships involving dysbiosis-linked taxa. Collectively, the correlation network elucidates a coordinated ecological reorganization towards a pro-inflammatory and pro-atherogenic microbial configuration in LDLr⁻/⁻+MPLs mice **(Fig. S4E)**.

### Functional prediction analysis reveals the roles of nanoplastics in atherosclerosis

Level 2 and Level 3 KEGG analysis done through PICRUSt2 demonstrated reprogramming of the gut microbiome in the LDLr⁻/⁻MPL group relative to both LDLr⁻/⁻ and C57BL/6 groups (KEGG level 2 and KEGG LEVEL 3). Nanoplastics exposure in the LDLr⁻/⁻+MPLs group enriched pathways as per KEGG level 2 that clearly showed that they were involved in cardiovascular diseases, the immune system, and others (**Fig. 7A**). At KEGG Level 3, we observed nanoplastics-induced enrichment of pathways linked to mucosal damage, oxidative stress, and inflammation, including epithelial stress, apoptosis, base excision DNA repair, and lipid metabolism. In contrast, pathways that can reduce inflammation and, hence, atherogenesis, such as xenobiotic processing, were comparatively reduced in the LDLr⁻/⁻+MPLs group (**Fig. 7 B).**

**Fig. 7.**
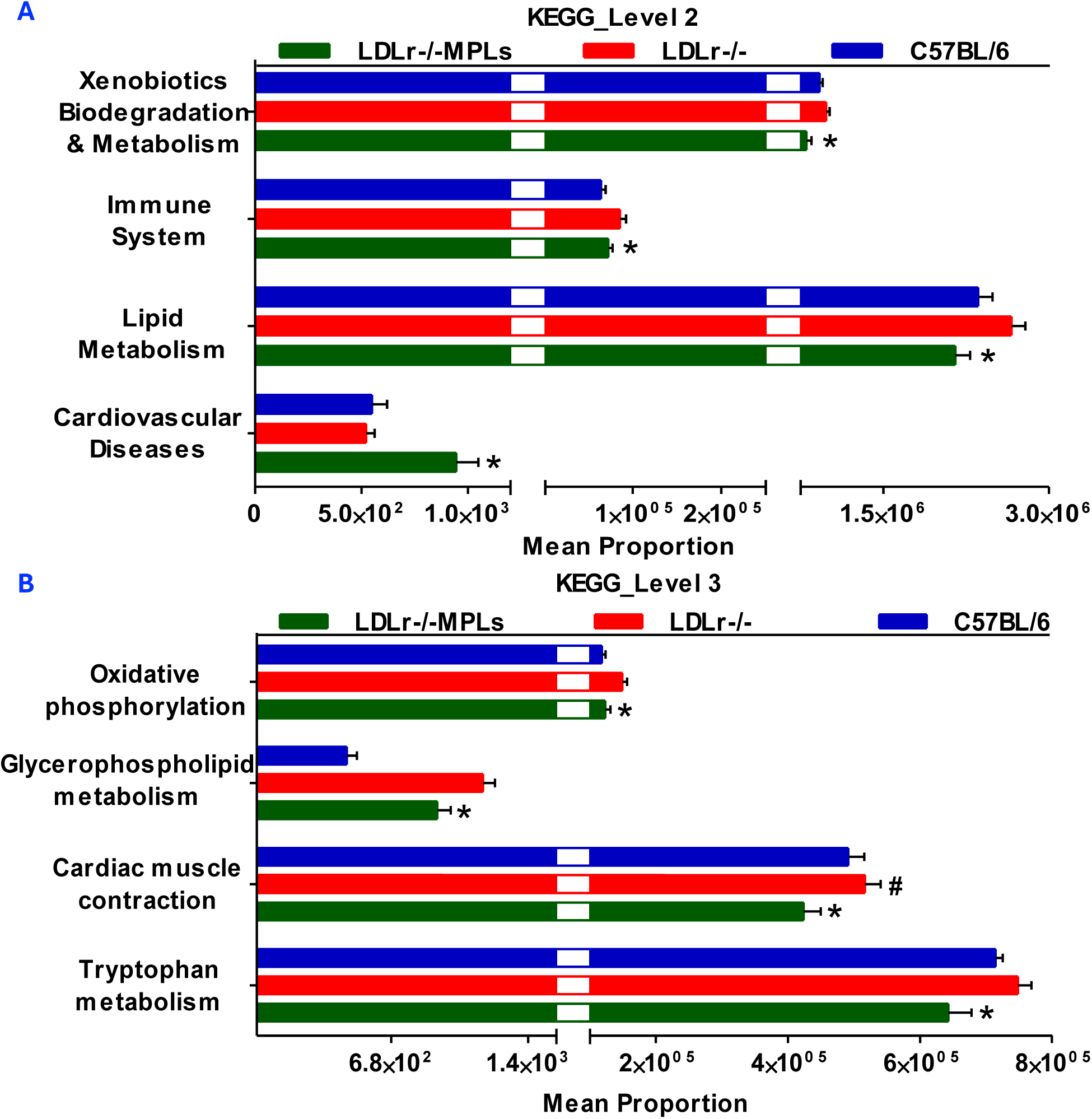
Predicted microbial functional pathways associated with microplastic exposure in LDLr⁻/⁻ mice. (A) KEGG Level 2 pathway analysis showing increased representation of microbial functional categories related to processes and diseases associated with atherosclerosis in LDLr⁻/⁻ mouse + MPLs mice compared with LDLr⁻/⁻ controls. (B) KEGG Level 3 pathway analysis demonstrating enrichment of specific predicted pathways linked to atherosclerosis-related processes in LDLr⁻/⁻+MPLs mice. * indicates a significant difference (p < 0.05) between LDLr⁻/⁻+MPLs and LDLr⁻/⁻ groups; # indicates a significant difference (p < 0.05) between LDLr⁻/⁻ and C57BL/6 mouse groups.

### Nanoplastics-induced gut dysbiosis in LDLr⁻/⁻ mice converges with human atherosclerosis

Comparative metagenomics analysis of nanoplastics-treated mice with human atherosclerosis metagenomics identified 18 shared genera whose abundance was dysregulated in both the LDLr⁻/⁻+MPLs mice group and human ACVD groups. Of the 154 unique harmonized genera found in both datasets (106 mouse-exclusive, 30 human-exclusive, and 18 shared), this overlap accounts for 11.7% (**Fig. 8A**). The shared genera include both taxa enriched in ACVD (*Actinomyces, Lactobacillus, Lachnospiraceae NK4A136 group, Rothia, Erysipelatoclostridium*, and others) and taxa depleted in ACVD (*Bacteroides, Eubacterium, Roseburia, Lachnospira*, and members of the *Lachnospiraceae* family) **(Fig. S6A)**. 6 out of the 18 shared genera exhibit concordant directional alterations, indicating the same direction of dysregulation in both mouse and ACVD (**Fig. 8B**). Four genera (*Rothia, Actinomyces, Lactobacillus, and Lachnospiraceae NK4A136 group*) were enriched in both LDLr⁻/⁻+MPLs mice and human ACVD patients (**Fig. 8B**). Two genera (*Bacteroides and Eubacterium*) were depleted in both (**Fig. 8B**). The remaining 12 genera showed discordant directional changes (depleted LDLr⁻/⁻+MPLs group but enriched in human ACVD) (**Fig. 8B**). Pathway annotation of the 18 shared identified four active mechanistic pathways (gut barrier integrity, LPS/endotoxin inflammation, SCFA/Butyrate synthesis, TMAO production) i.e SCFA/butyrate synthesis (10 genera, all scored −1 as depleted protective producers), gut barrier integrity (8 genera; 5 protective, 3 pro-atherogenic), LPS/endotoxin (5 genera; 4 pro-atherogenic, 1 protective), and TMAO production (2 genera, both pro-atherogenic); *Tyzzerella* and *Lachnospiraceae_unclassified* carried no assignable pathway score **(Fig. S6B).** Cardiovascular disease (CVD) evidence mapping across the same genera produced 41 supported genus-CVD associations out of 90 possible cells (11 strong, 30 Good), with ACVD showing the highest genus coverage (14/18; **Fig. 8C**). SCFA/butyrate depletion and gut barrier integrity each had genus-level evidence across all five CVD categories. At the same time, LPS/endotoxin and TMAO pathways were represented in four and three CVD categories, respectively (**Fig. 8D**).

**Fig. 8.**
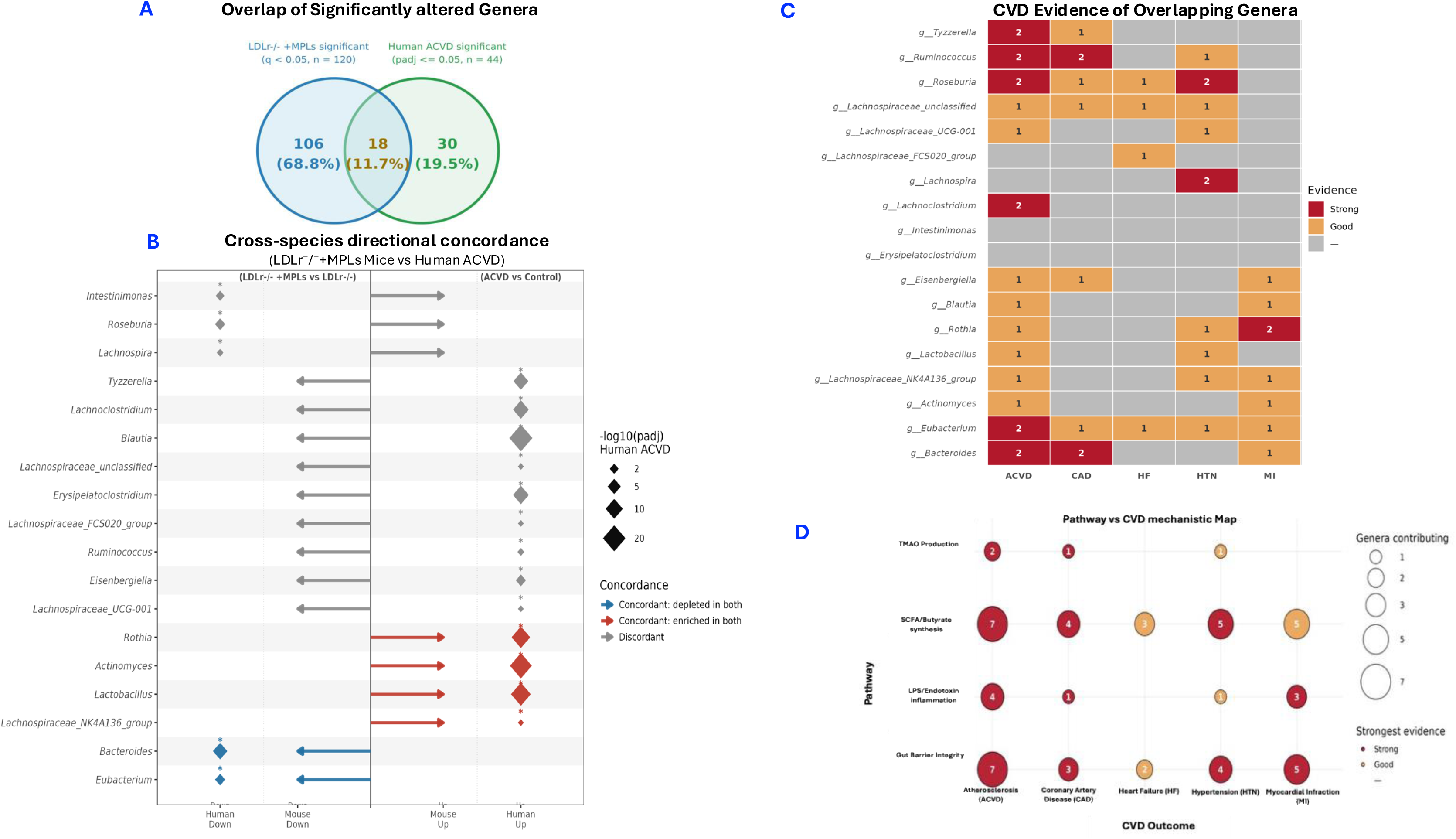
Nanoplastics-induced gut dysbiosis in LDLr⁻/⁻ mice converges with human atherosclerosis and cardiovascular disease pathways. (A) Venn diagram of the overlap between gut genera significantly altered in LDLr⁻/⁻+NPLs mice versus LDLr⁻/⁻ controls (padj ≤ 0.05; n = 120) and in human ACVD patients versus healthy controls (padj ≤ 0.05; n = 44), with 18 shared genera (11.7%). (B) Directional concordance for the 18 shared genera. Arrows (left): direction of change in LDLr⁻/⁻+NPLs versus LDLr⁻/⁻ (rightward = enriched, leftward = depleted); diamonds (right): direction and significance in human ACVD versus controls (Jie et al., 2017), sized by −log₁₀(padj), with * marking padj ≤ 0.05. (C) Evidence heatmap linking the 18 shared genera to five cardiovascular disease categories (atherosclerosis/ACVD, coronary artery disease, heart failure, hypertension, myocardial infarction); tile color indicates strong, good, or no evidence. (D) Bubble plot integrating four mechanistic pathways (gut barrier, LPS, SCFA, TMAO) with the five CVD categories; bubble size indicates the number of shared genera linking each pathway to each disease, and color indicates the strongest evidence level for that combination.

### Hepatic transcriptome and gut microbiome integrative analysis reveals that nanoplastics exposure suppresses a hepatic xenobiotic detoxification module linked to gut microbial shift in LDLr⁻/⁻ mice

Tripartite network linked hepatic co-expression module with their most strongly varying gut bacterial genera, hence demonstrating a coordinated gut-liver response to nanoplastics exposure in LDLr ⁻/⁻ mice (**Fig. 9A**). Four hepatic DEGs modules (M1 with 61 genes, M2 with 25 genes, M3 with 7 genes, and M4 with 6 genes) were found through hierarchical clustering out of which two differed significantly in eigengene activity between exposure groups: M1, in which 54 of 61 genes (89%) were downregulated, showed reduced activity, and M2, comprising 25 uniformly upregulated genes, showed elevated activity in nanoplastics-exposed mice (both p < 0.001, Welch’s t-test) (**Fig. 9B**). Enrichment analysis showed that M1 is a hepatic xenobiotic detoxification program covering phase I (cytochrome P450) and phase II (glucuronidation) pathways (top GO and KEGG terms respectively, with hub genes including *Pgk1, Tmco1, S1pr1, Enpp5, and Tmed2b*, all with kME ≥ 0.993), while M2 was enriched for cell cycle control terms (regulation of DNA replication) (**Fig. 9C**). The network showed a coordinated disruption of the hepatic transcriptome and gut microbiota consistent with microplastic driven perturbation of the gut liver axis.

**Figure 9.**
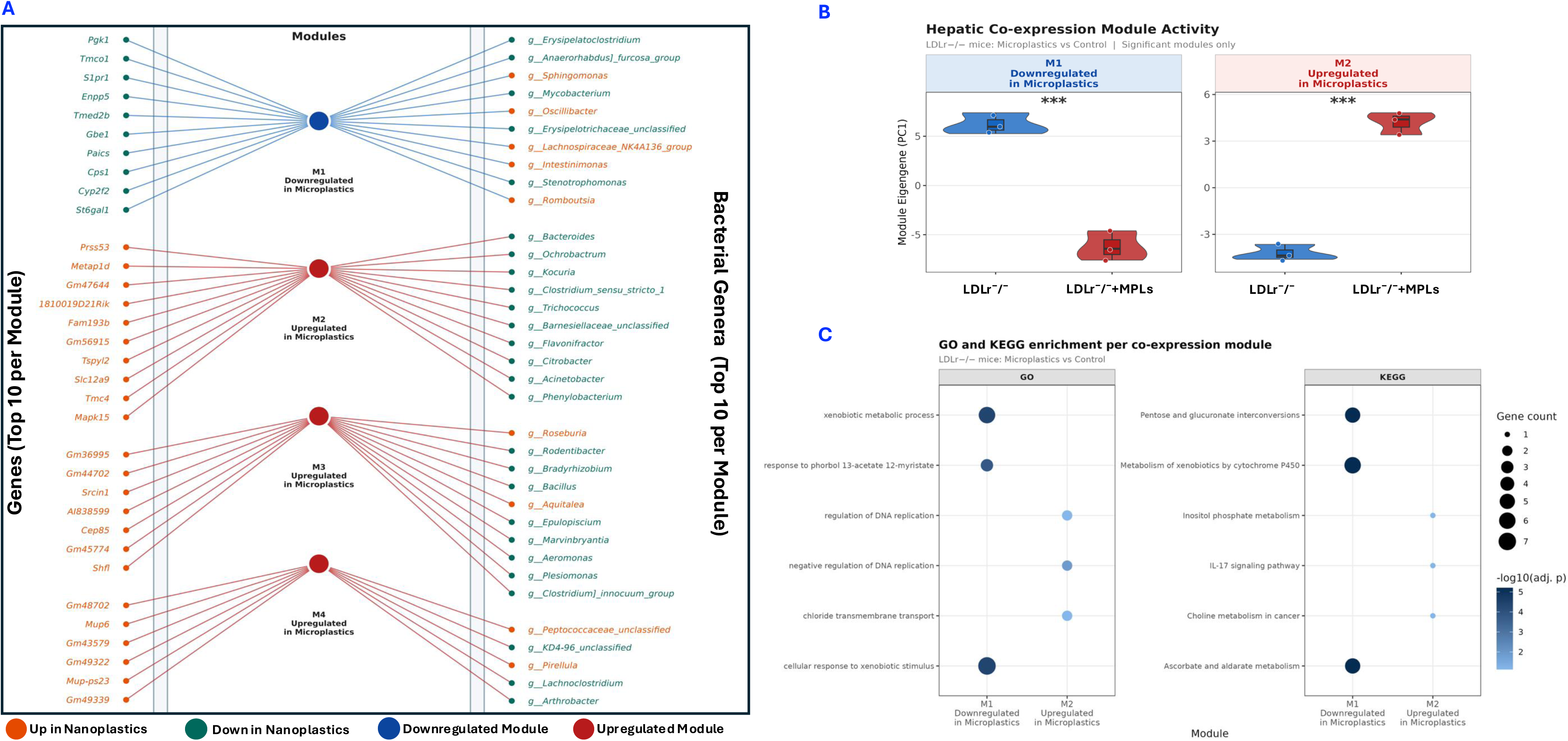
Integrative hepatic transcriptomic and gut microbiome analysis of nanoplastics exposure in LDLr⁻/⁻ mice. (A) left: Tripartite network linking hepatic DEGs, center: co-expression modules, and right: differentially abundant gut bacterial genera. Edges connect genes to their assigned module and modules to their associated genera, colored by module. Node color indicates direction of change in nanoplastics-exposed versus control mice (orange = upregulated/enriched; teal = downregulated/depleted); module nodes are colored by dominant member-gene direction (blue = M1, predominantly downregulated; red = M2–M4, predominantly upregulated). The top 10 hub genes (by kME) and top 10 differentially abundant genera (by absolute bicor to the module eigengene) are shown per module. (B) Module eigengene activity for significantly altered modules. Violin plots (with overlaid boxplots and data points) show eigengene values for M1 and M2 in control (blue, n = 3) and nanoplastics-exposed (red, n = 3) mice. M1 (61 genes, 54 downregulated [89%]) and M2 (25 genes, all upregulated) differed significantly (Welch’s t-test, ***p < 0.001); M3 and M4 were not significant and are not shown. (C) GO Biological Process and KEGG enrichment of member genes from M1 (downregulated) and M2 (upregulated). The top three terms per module per database are shown; dot size reflects annotated gene count and color intensity reflects −log₁₀(adjusted p-value). M2 KEGG terms with ≤ 2 genes should be interpreted cautiously given the small module size.

### Metabolomic profiling reveals distinct metabolic reprogramming following nanoplastics exposure

The three groups demonstrated clear separation as shown in PCA of the filtered feature matrix with PC1 and PC2 accounting for 40.0% and 20.1%, respectively (**Fig. 10A**). One-way ANOVA identified a set of metabolic features that had significantly different abundance in all three groups. Hierarchical clustering further resulted in four interpretable feature clusters from the top 41 ANOVA-significant features using Ward’s D2 linkage on group mean z-score (**Fig. 10B**). The first cluster comprised lipid species (LysoPE(20:5), LysoPC(O-18:0), MD920:3), PC(22;6), the vitamin A metabolite (9-cis-retinol), and ophthalmic acid (an oxidative stress-response glutathione analog) that were elevated in C57BL/6 but reduced in both LDLr⁻/⁻ and LDLr⁻/⁻+MPLs groups. The second cluster contained features such as N-oleoylethanolamine, eicosatrienoic acid, and porphobilinogen that were more enriched in LDLr⁻/⁻+MPLs group as compared to the LDLr⁻/⁻ and C57BL/6 group. A third cluster showed features such as phenylalanyl-asparagine, benzoic acid, coumaric acid, 2-acetyl-4-methylpyridine, and pyroglutamic acid, which were enriched only in the LDLr⁻/⁻+MPLs group. A small set of features was depleted in both LDLr⁻/⁻ and LDLr⁻/⁻+MPLs groups as compared to the C57BL/6 group. Therefore, the heatmap demonstrates that exposure to nanoplastics increased a subset of disease-related features (cluster 2) and that a specific set of features emerged only in the nanoplastics-exposed group (cluster 3), suggesting that nanoplastics create a metabolic environment that affects the disease state. To determine metabolic effects nanoplastics alone, LDLr⁻/⁻+MPLs was compared to LDLr⁻/⁻, which yielded 27 statistically significant feature–annotation pairs (p < 0.05), and 10 pairs after deduplication by m/z-retention having a role in atherosclerosis including Neurine, Gamma-Glutamylcysteine, Phenylalanyl-Asparagine, all-trans-3,4-Didehydroretinoate, Hallacridone, Glycodiazine, 6-O-Desmethyldonepezil, Ambenonium, Maltohexaose (**Fig. 10C**). Gamma-glutamylcysteine which is a rate-limiting precursor glutathione was significantly reduced in LDLr⁻/⁻+MPLs group compared to both LDLr⁻/⁻ and C57BL/6. Neurine, a choline-derived trimethylamine compound, was similarly depleted in LDLr⁻/⁻+MPLs, suggesting altered flux through the choline–trimethylamine metabolic axis. In contrast, phenylalanyl-asparagine, all-trans-3,4-didehydroretinoate, 6-O-desmethyldonepezil, ambenonium, and maltohexaose were substantially increased in the LDLr⁻/⁻+MPLs group (**Fig. 10C**). Collectively, these results suggest that nanoplastics exposure produces measurable and statistically significant changes in hepatic metabolome profiles of mice.

**Fig. 10. Untargeted.**
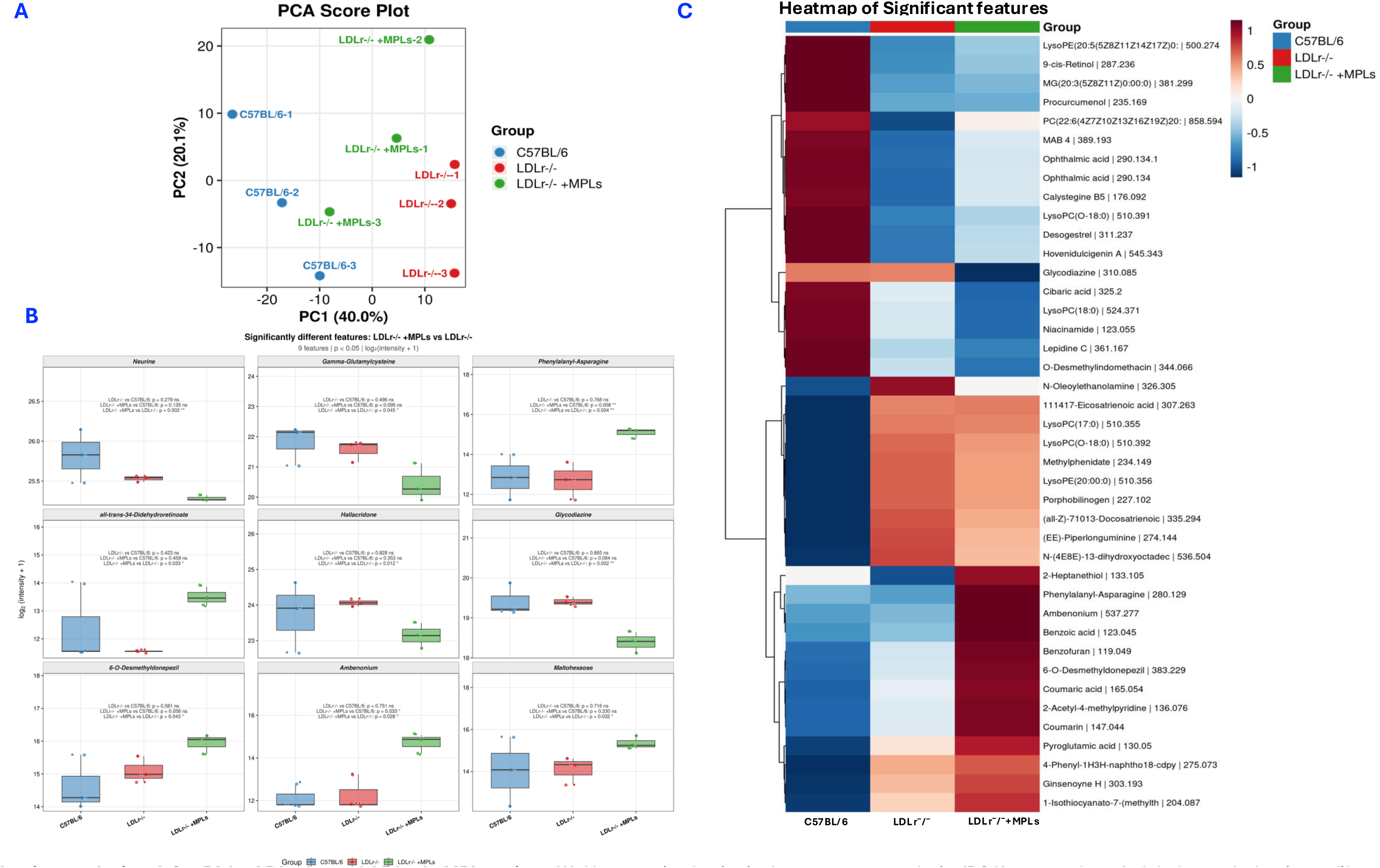
m**e**tabolomics **analysis of C57BL/6, LDLr⁻/⁻, and LDLr⁻/⁻+MPLs mice.** (A) Unsupervised principal component analysis (PCA) score plot of global metabolomic profiles across the three experimental groups: C57BL/6 mouse (blue), LDLr⁻/⁻ mouse (red), and LDLr⁻/⁻+MPLs (green). Each point represents an individual biological replicate. (B) Hierarchical clustering heatmap of the 41 metabolic features that reached statistical significance by one-way ANOVA (p < 0.05). Values represent group mean intensities normalized as log₂-transformed z-scores. Rows were clustered using Ward’s D2 linkage, and columns were arranged in biological group order. Red and blue indicate relatively higher and lower feature abundance, respectively. (C) Boxplots showing the distribution of log₂-transformed intensities for the 9 metabolites that differed significantly between LDLr⁻/⁻+MPLs and LDLr⁻/⁻ groups (p < 0.05). Individual data points are overlaid for each group. P-values for all three pairwise comparisons are annotated within each panel, with significance indicated as follows: *p < 0.05, **p < 0.01, ***p < 0.001; ns, not significant.

## Discussion

Microplastics and nanoplastics (M-NPLs) are ubiquitous environmental contaminants that enter the human body through food, drinking water, and inhalation and accumulate in various tissues, including blood, the placenta, the lung, the gut, and, most importantly, atherosclerotic plaque (Schwabl, Koppel et al. 2019, Amato-Lourenco, Carvalho-Oliveira et al. 2021, Ragusa, Svelato et al. 2021, Leslie, van Velzen et al. 2022, Khan and Jia 2023, Marfella, Prattichizzo et al. 2024). Increasing evidence indicates that M-NPLs exposure is associated with cardiovascular disease (van Grevenynghe, Monteiro et al. 2006, Oesterling, Toborek et al. 2008, Hu and Palić 2020, Lett, Hall et al. 2021, Marfella, Prattichizzo et al. 2024). Despite these findings, the underlying mechanisms of M-NPLs exposure in the development of atherosclerosis remain unclear and require further investigation. To the best of our knowledge, this is the first comprehensive multi-omics investigation of the atherogenic effects of 80 nm polystyrene nanoplastics in LDLr⁻/⁻ mice. Previous studies have predominantly utilized ApoE⁻/⁻ mice to investigate the cardiovascular effects of nanoplastics exposure, whereas studies in LDLr⁻/⁻ mice remain limited. In particular, the atherogenic effects and underlying molecular mechanisms of chronic nanoplastic exposure in LDLr⁻/⁻ mice have not been comprehensively characterized. To address this gap, we performed a comprehensive multi-omics investigation of chronic 80 nm polystyrene nanoplastic exposure in LDLr⁻/⁻ mice, integrating aortic plaque assessment with hepatic transcriptomics, genome-wide alternative splicing analysis, gut microbiome profiling, and untargeted metabolomics to provide a systems-level understanding of the biological mechanisms linking chronic nanoplastic exposure to atherosclerosis. Importantly, cross-validation of exposure-associated molecular signatures with human atherosclerotic datasets enhances the translational relevance of our findings. This integrative approach enables the identification of coordinated alterations across vascular, metabolic, and microbial pathways. It provides novel evidence that nanoplastic exposure may promote atherosclerosis through interconnected host–microbiome–metabolic networks.

Using LDLr⁻/⁻ mice fed an atherogenic diet as a model, this study demonstrated that long-term exposure to nanoplastics accelerates the progression of atherosclerosis. Our assessments of vascular lesions, specifically quantitative analysis using Oil Red O staining, revealed a significant increase in lipid deposition within the aortic intima across all groups exposed to polystyrene nanoplastics. At the vascular level, a significant increase in aortic intimal lipid deposition suggests the involvement of multiple synergistic mechanisms, such as lipid metabolic dysregulation, impaired clearance, endothelial injury, and immune cell infiltration (Libby 2002, Libby, Ridker et al. 2002, Gimbrone and García-Cardeña 2016). These findings are consistent with previous mouse studies: oral ingestion of 1 µm microplastics induces atherosclerotic lesions in LDLr⁻/⁻ models (Lin, Pan et al. 2025), while long-term exposure to 50 nm polystyrene nanoplastics in ApoE⁻/⁻ mice promotes plaque formation, aortic stiffening, and macrophage M1 polarization by upregulating MARCO expression and disrupting lipid metabolism (Wang, Liang et al. 2023). Our findings help explain the results of a recent prospective cohort study that detected unusually high concentrations of nano- and micro-scale plastics (averaging approximately 27 µg/mg of plaque tissue) in human carotid atherosclerotic plaques (Marfella, Prattichizzo et al. 2024). Notably, the presence of these plastics in plaques was significantly associated with a 4.53-fold increased risk of major adverse cardiovascular events (Marfella, Prattichizzo et al. 2024); this not only suggests for the first time in humans that the accumulation of plastic particles may be a potential risk factor for cardiovascular disease but also underscores the urgency of elucidating the mechanisms by which they promote atherosclerosis.

Using a multi-omics approach, integrating transcriptomics, alternative splicing, gut metagenomics, and metabolomics, and comparing the results with human atherosclerosis data, this study reveals that the acceleration of atherosclerosis by nanoplastics is not mediated by a single pathway but results from the synergistic interplay of dysregulated hepatic gene expression, systemic metabolic remodeling, and gut microbiota imbalance.

The hepatic transcriptomic profile provided mechanistic insight into the roles of chronic exposure to nanoplastics in atherosclerosis. It was far more pronounced at the transcript level than at the gene level: 2,297 differentially expressed transcripts (DETs) versus 1,183 differentially expressed genes (DEGs), with 82% of all significant loci detectable only at the transcript level. Since transcript-level changes can shape protein structure, localization, stability, and signaling, these regulatory layers carry information that gene-level analysis could not identify, a gap that is particularly important for immune and chronic and inflammatory diseases (Kim, Pham et al. 2018, Tao, Zhang et al. 2024). Alternative splicing analysis further supported this, identifying 1321 significant splicing events dominated by intron retention (IR, 41.56%) and skipped exon (SE, 22.79%). With recent advances in alternative splicing, intron retention is now considered a regulated process rather than a random splicing error. It affects protein production by retaining transcripts in the nucleus, decreasing RNA Stability, triggering nonsense-mediated decay, often in response to cellular stress or differentiation signals (Porras-Tobias, Caldera et al. 2025). Skipped exons also modify transcriptomic outcomes by generating isoforms. The interpretation that chronic exposure to polystyrene nanoplastics alters hepatic RNA processing in a manner consistent with endoplasmic reticulum and proteostatic stress is supported by gene ontology enrichment for endoplasmic reticulum processes, oxidoreductase activity, lipopolysaccharide responses, and xenobiotic and detoxification pathways, including cytochrome P450 and glucuronidation. Unlike previous micro- and nanoplastics toxicity studies (Lu, Wan et al. 2018, Chen, Zhuang et al. 2022, Lee, Lin et al. 2024, Zhao, Adiele et al. 2024), our data show that polystyrene nanoplastics exposure affects hepatic RNA processing. This mechanism has not previously been implicated in microplastic and nanoplastics toxicity studies.

Notably, this study found that nanoplastics exposure induces alternative splicing and isoform switching as a previously unrecognized regulatory layer through which nanoplastics exposure impacts atherosclerosis-related genes. Specifically, 1-acylglycerol-3-phosphate O-acyltransferase 3 (*Agpat3*), which is involved in lipid metabolism; dipeptidyl peptidase 9 (*Dpp9*), which regulates immune and inflammatory pathways; and eukaryotic translation initiation factor 4A2 (*Eif4a2*), which controls cap-dependent translation, all exhibited significant changes in transcript structure following nanoplastics exposure; these alterations may remodel metabolic and immune signaling. This finding aligns with an emerging body of work demonstrating that alternative splicing is a pervasive and disease-relevant regulatory layer in the cardiovascular system (van den Hoogenhof, Pinto et al. 2016, Huynh 2024), with intron retention specifically now recognized as one of the most frequent AS events in ischemic cardiomyopathy (Cao, Wei et al. 2024) and as a feature of inflammatory NF-κB signaling, in which alternative splicing generates functionally distinct protein isoforms (Leeman and Gilmore 2008), consistent with this, inflammation-induced alternative splicing in human endothelial cells produces novel isoforms of cardiovascular-disease-associated genes (Golebiewski, Stolze et al. 2025). Our isoform-switching analysis further identified 153 genes in which transcripts from the same locus moved in opposite directions, indicating genuine regulatory switching rather than uniform up- or down-regulation. The concordance between isoform-switch bar plots and splice-junction usage in sashimi plots (for *Ubl7, Canx, Sipa1, Cbs, and Chka*) provides the strongest evidence that these events are biologically real. At the pathway level, enrichment of fatty-acid and peroxisomal programs implicates altered lipid oxidation and trafficking. In contrast, enrichment of bile-secretion and xenobiotic terms points to disrupted hepatic transport and detoxification. These signals converge on a focused atherosclerosis-gene set, in which 52 curated genes were dysregulated with a striking 3.3:1 downregulation bias: coordinate suppression of cholesterol biosynthesis (*Cyp51, Sqle, Idi1*), reverse cholesterol transport (*Abca1, Ldlr*), mitochondrial function (*Sdhd, Prdx3*), and cardioprotective lipid-mediator synthesis (*Cyp2j5*), set against upregulation of the ER-stress marker *Ddit3*, the pro-apoptotic factor *Bax*, and mitochondrial complex subunits (*Ndufa3, Ndufb7, Uqcrq*). This predominantly suppressive rewiring of atheroprotective programs mirrors the plaque burden quantified at the aortic root and is consistent with classical models in which lipid handling, endothelial dysfunction, and inflammation jointly drive atherogenesis (Libby, Ridker et al. 2002, Tabas, García-Cardeña et al. 2015, Gimbrone and García-Cardeña 2016), positioning the liver as a central coordinator of the systemic atherogenic response to nanoplastics exposure.

To explore the correlation between our microplastic animal model experimental results and the clinical manifestations and progression of human atherosclerosis, we compared the liver transcriptomes of mice exposed to polystyrene nanoplastics with a human peripheral atherosclerosis dataset (GSE100927). Notably, 183 overlapping genes were identified, exhibiting highly conserved regulatory patterns across both species; specifically, macrophage- and inflammation-related genes, such as *Trem2*, *Cd68*, *Tyrobp*, *Lpl*, *Cyth4*, *Pla2g7*, *Vav1*, and *Capg*, were consistently upregulated and collectively enriched in pathways related to immune responses, lipid metabolism, and atherosclerosis. This cross-species transcriptomic conservation holds threefold translational significance: first, it demonstrates that the microplastic-induced molecular alterations observed in mouse livers are not species-specific phenomena but are biologically relevant within the context of human atherosclerotic pathology; second, it aligns with prior clinical The depletion of γ-glutamylcysteine observed in this study aligns with the phenotype of accelerated atherosclerosis, suggesting an insufficient supply of glutathione precursors and compromised hepatic antioxidant defensestudies, which have shown that the detection of micro-and nanoplastics within plaques is significantly associated with elevated inflammatory markers and an increased risk of cardiovascular events (Marfella, Prattichizzo et al. 2024, Prattichizzo, Ceriello et al. 2024, Dowling 2025), thereby corroborating our findings from an independent perspective; and third, and most importantly, this study establishes, for the first time at the transcriptomic level, a direct molecular link between nanoplastics exposure, hepatic responses, and human atherosclerosis, providing a robust theoretical and experimental foundation for the future screening of biomarkers and the development of intervention strategies.

Given that micro- and nanoplastics have been detected in the human gut (Schwabl, Koppel et al. 2019), that the pathogenic role of exogenous substance-induced gut microbiota dysbiosis in cardiovascular disease is well-established (Garcia-Rios, Torres-Peña et al. 2017, Kitai and Tang 2018, Witkowski, Weeks et al. 2020), and that orally ingested particles first interact with the host at the intestinal surface (Schwabl, Koppel et al. 2019, Roslan, Lee et al. 2024, Zeng, Li et al. 2024), investigating the gut microbiota’s response to nanoplastics exposure holds significant mechanistic importance. Our data indicate that nanoplastics exposure does not significantly alter overall within-sample (alpha diversity) between between the diet- and genotype-matched LDLr⁻/⁻+MPLs and LDLr⁻/⁻ groups, but instead drives dysbiosis by selectively remodeling community structure (resulting in significant changes in beta diversity) (Jonsson and Bäckhed 2017, Tang, Kitai et al. 2017). The compositional change in the microplastic-induced gut microbiome was directionally similar to that observed in cardiovascular risk. The enrichment of *Streptococcus* is significant due to its correlation with coronary artery calcium and subclinical atherosclerosis, as well as its presence in plaque, oral, and gut regions of cardiovascular patients (Koren, Spor et al. 2011, Jie, Xia et al. 2017, Pisano, Bugli et al. 2023). In addition, nanoplastics exposure increased *Faecalibaculum* abundance, a finding consistent with the proliferation of this genus observed in cardiovascular risk models following exposure to the environmental contaminant DEHP (Chai, Wen et al. 2023). In contrast, genera that are beneficial for cardiovascular health, such as *Bacteroides* (which promotes immune tolerance by balancing Th1/Th17 responses (Round, Lee et al. 2011)) and *Christensenellaceae* (which are linked with reduced cardiometabolic risk, anti-inflammatory properties, and barrier integrity (Goodrich, Waters et al. 2014, Waters and Ley 2019, Okalin, Arslan et al. 2025)), were depleted by nanoplastics exposure. Additionally, our functional prediction (PICRUSt2) demonstrated enrichment of pathways associated with mucosal stress, oxidative stress, apoptosis, and inflammation, along with a relative decline in xenobiotic processing, reflecting hepatic transcriptomic changes and suggesting a coordinated gut-liver response.

While our microbiome data demonstrate that nanoplastics exposure alters gut microbiome dysbiosis in a pro-atherogenic manner, it is still not clear whether this alteration corresponds to human cardiovascular disease. To address this translation gap, we used a novel approach for the first time to systematically compare our polystyrene nanoplastics metagenomics data with the well-characterized human ACVD metagenome from the Jie et al. study (Jie, Xia et al. 2017). Among 154 harmonized genera, 18 showed dysregulation in the groups (polystyrene nanoplastics-treated mice and human ACVD patients). Six demonstrated directional concordance, comprising four genera enriched in both contexts (*Rothia, Actinomyces, Lactobacillus,* and the *Lachnospiraceae NK4A136 group*) and two genera depleted in both (*Bacteroides* and *Eubacterium*). Pathway scoring identified that SCFA/butyrate synthesis serves as a mechanistic hub, with 10 out of eighteen genera identified as SCFA producers, whose reduction is pro-atherogenic. This aligns with butyrate’s cardioprotective function in facilitating regulatory T-cell differentiation, cholesterol efflux, and tight-junction integrity, as well as the consistent decline of producers like *Roseburia* across human cohorts (Jie, Xia et al. 2017). The concordant enrichment (both in polystyrene nanoplastics-treated mice and in human ACVD) of *Actinomyces* and *Rothia* (oral-derived pathobionts) suggests a second pathway to vascular injury: pathobionts crossing the microplastic-compromised intestinal barrier, releasing lipopolysaccharide into the bloodstream, and triggering systemic inflammation (Huang, Weng et al. 2021, Zeng, Li et al. 2024). Together, commensal depletion and pathobiont enrichment therefore suggest at least two parallel microbial axes, i.e., metabolic and immunological, in microplastic-driven atherogenesis. The enrichment of *Ruminococcus* and *Lachnoclostridium* suggests a TMAO production axis, possibly driven by SCFA depletion that makes the gut environment more hospitable to TMAO-producing bacteria. TMAO is a well-known cardiovascular risk factor that promotes vascular inflammation and is associated with adverse clinical outcomes (heart attacks and strokes) (Wang, Klipfell et al. 2011, Seldin, Meng et al. 2016, Heianza, Ma et al. 2017). To our knowledge, this is the first polystyrene nanoplastics exposure study to perform cross-species validation at two independent molecular levels (hepatic transcriptomics and gut metagenome), while comparing the experimental findings to human atherosclerotic disease.

The gut-liver axis is a well-recognized regulator of metabolic and inflammatory disease; however, integrated studies linking microplastic-associated hepatic transcriptomes with gut microbiomes remain lacking. Given our previous finding that microplastic-induced gut microbiome signatures align with the metagenomic profiles associated with human atherosclerotic cardiovascular disease (ACVD), this study further investigates the synergistic interplay between the hepatic transcriptome and the gut metagenome in the context of atherosclerosis. Our studies indicated that two hepatic modules were significantly different in eigengene activity: M1, a large, predominantly downregulated module (61 genes, 89% suppressed), and M2, an upregulated module (25 genes). M1 was prominently enriched for xenobiotic-detoxification pathways, including phase I cytochrome P450 metabolism and phase II glucuronidation, with hub genes such as *Pgk1* and the atheroprotective lipid-signaling/endothelial-barrier mediator *S1pr1* (Zanger and Schwab 2013, Blaho and Hla 2014). Its coordinated suppression indicates that nanoplastics exposure may weaken hepatic detoxification processes, hence potentially limiting the liver’s ability to metabolize xenobiotic stressors. This finding is consistent with reports of microplastics-induced disruption of hepatic detoxification through gut–liver signaling (Wang, Deng et al. 2024) and extends those observations to a network-level response in an atherosclerosis-prone model. The enrichment of M2 for cell cycle and DNA replication is consistent with microplastics-induced hepatocyte stress and impaired regenerative capacity (Tripathi, Debelius et al. 2018). Notably, the significant hepatic modules were linked to a distinct and biologically coherent set of gut microbial genera. Suppression of the M1 detoxification module co-occurred with depletion of *Bacteroides, Erysipelatoclostridium*, and other SCFA-associated commensals, which were similar to studies reporting that micro- and nanoplastics reduce *Bacteroidetes* and SCFA-producing taxa (Huang, Weng et al. 2021). The loss of these microbial signals may eliminate an important regulatory input affecting hepatic lipid metabolism, inflammatory tone, and detoxification-related gene expression (Brandl, Kumar et al. 2017, Tripathi, Debelius et al. 2018, Albillos, De Gottardi et al. 2020, Tilg, Adolph et al. 2022). Since both datasets (hepatic transcriptome and gut microbiome) are from the same mice (same study), these associations reveal a strong biological linkage. This, to our knowledge, is also the first study of its kind to integrate the hepatic transcriptome with the gut microbiome in response to nanoplastics exposure and suggests that hepatic and microbial remodeling are coordinated responses that could lead to atherosclerosis.

Our research indicates that nanoplastics exposure remodels the hepatic transcriptional profile and alters the structure of gut microbial communities. To investigate whether these upstream transcriptional changes translate into biochemical disturbances at the metabolic level, we conducted untargeted hepatic metabolomic analysis. Principal component analysis revealed distinct separation among the three experimental groups; notably, the metabolite profile of the LDLr⁻/⁻ + MPLs group differed significantly from that of the control group, with the significantly altered metabolites primarily enriched in three pathways closely associated with cardiovascular disease. Of particular note, levels of γ-glutamylcysteine, the direct precursor to glutathione and the product of its rate-limiting synthesis enzyme, decreased significantly following nanoplastics exposure. Previous studies have shown that vascular glutathione levels decline during the early stages of atherosclerosis; oxidized low-density lipoprotein (ox-LDL) can induce glutamate-cysteine ligase (GCL), the rate-limiting enzyme of glutathione synthesis, and intracellular glutathione synthesis in macrophages influences lesion progression (Bea, Hudson et al. 2003, Biswas, Newby et al. 2005, Callegari, Liu et al. 2011). Furthermore, polystyrene micro- and nanoplastics have been shown to directly elevate hepatic reactive oxygen species (ROS) levels and inhibit antioxidant enzyme activity (Zou, Qu et al. 2023). The depletion of γ-glutamylcysteine observed in this study suggests that, under chronic nanoplastic-induced oxidative stress, consumption of glutathione precursors outpaces any compensatory upregulation of GCL, resulting in compromised hepatic antioxidant defense consistent with the phenotype of accelerated atherosclerosis; these findings are consistent with the known properties of polystyrene micro- and nanoplastics regarding ROS elevation and antioxidant enzyme inhibition, as well as the early glutathione depletion observed before and during the progression of atherosclerosis (Bea, Hudson et al. 2003, Biswas, Newby et al. 2005, Callegari, Liu et al. 2011). We found that neurine (a quaternary amine derived from choline) was also reduced following MPLs treatment, which aligns with disruption of gut choline metabolism that favors the pro-atherogenic trimethylamine/TMAO pathway (Wang, Klipfell et al. 2011, Seldin, Meng et al. 2016, Heianza, Ma et al. 2017). The increase in all-trans-3,4-didehydroretinoate, together with the depletion of 9-cis-retinol, demonstrates that MPLs exposure may disrupt retinoid signaling, which regulates macrophage cholesterol efflux and lesion formation (Cassim Bawa, Gopoju et al. 2022), suggesting oxidative stress on hepatic retinoid capacity rather than beneficial retinoid activation. Collectively, the synergistic effects of oxidative stress, inflammatory responses, activation of the TMAO pathway, and disordered cholesterol metabolism, all induced by nanoplastics exposure, collectively promote lipid deposition in the arterial wall, ultimately accelerating the formation of atherosclerotic plaques.

In summary, this study demonstrates that chronic polystyrene nanoplastics exacerbate atherosclerotic lesions in LDLr⁻/⁻ mice and is associated with coordinated remodeling transcriptomic, metabolomic, and metagenomic. Notably, beyond metabolic and microbial alterations, post-transcriptional regulatory events, such as alternative splicing, intron retention, exon skipping, and isoform switching, also play pivotal roles in the response to nanoplastics. Through multi-omics integrative analysis (with a particular focus on the transcriptomic level), this study offers new insights into how nanoplastics exposure exacerbates cardiometabolic diseases. Furthermore, the identified isoform markers and host-microbiota interaction signatures may serve as early warning indicators and potential therapeutic targets for microplastic-associated cardiovascular diseases.

## Supporting information

https://uncg-my.sharepoint.com/:u:/g/personal/a_khan10_uncg_edu/IQDdOD-nqdMjS5LEbGvLDjUjAZGJiuZ5SVDtwyL_oY1JXp8?e=PeO5hi

## Acknowledgements

The authors thank the Triad Mass Spectrometry Facility in the Department of Chemistry and Biochemistry at the University of North Carolina at Greensboro for LC-MS metabolomics analysis. We gratefully acknowledge the use of the Longleaf high-performance computing cluster, provided by UNC Research Computing at the University of North Carolina at Chapel Hill, for the bioinformatics and data analyses in this study.

## CRediT authorship contribution statement

**Ajmal Khan:** Conceptualization, Methodology, Investigation, Formal analysis, Software, Data curation, Visualization, Writing – original draft. **Delicia Esther Cardenas Vasquez:** Investigation. **Yaru Si:** Investigation. **Warren S. Vidar:** Investigation. **Kerui Wu:** Review and Editing. **Norman Chiu:** Conceptualization, Methodology, Resources. **Zhenquan Jia:** Conceptualization, Methodology, Supervision, Project administration, Funding acquisition, Review and Editing.

## Declaration of Interest

All the authors declare that they have no competing interest that could influence the work reported in this paper

## Data availability statement

The datasets generated and analyzed in this study will be made publicly available upon publication.

## Notes

### Competing Interest Statement

The authors have declared no competing interest.

