## Supplementary material for "Chronic Polystyrene Nanoplastics Exposure Reprograms Gene Expression, Alternative Splicing, and Disrupts Host–Microbiome–Metabolic Networks to Promote Atherosclerosis in LDLr⁻/⁻ Mice": https://uncg-my.sharepoint.com/:u:/g/personal/a_khan10_uncg_edu/IQDdOD-nqdMjS5LEbGvLDjUjAZGJiuZ5SVDtwyL_oY1JXp8?e=PeO5hi: Supplementary Figures_Nanoplastics and Atherosclerosis.pdf

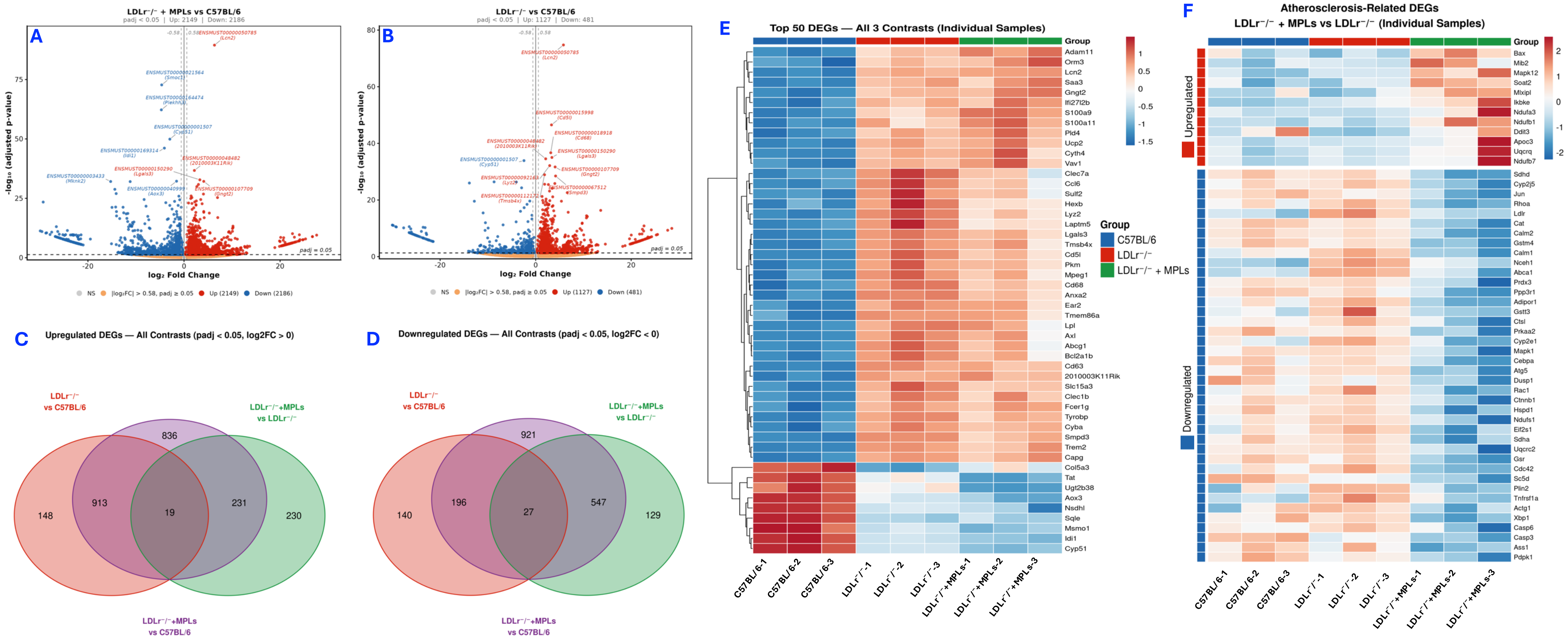


**Fig. S1. Extended gene-level evidence for microplastic-induced transcriptional reprogramming in LDLr$^{-/-}$ liver.** (A) Volcano plot of gene-level differential expression in the LDLr$^{-/-}$+MPLs vs C57BL/6 contrast (padj < 0.05). Red points indicate significantly upregulated genes, blue points significantly downregulated genes, grey points non-significant genes, and orange points genes with $|\log_2 FC| > 0.58$ but padj ≥ 0.05. The horizontal dashed line marks padj = 0.05 and vertical dashed lines mark $\log_2 FC = \pm 0.58$. The top 10 most significantly upregulated and downregulated genes are labeled. (B) Volcano plot of gene-level differential expression in the LDLr$^{-/-}$ vs C57BL/6 contrast (padj < 0.05), using the same color scheme and labeling criteria as panel A. (C) Three-way Venn diagram restricted to significantly upregulated DEGs ($\log_2 FC > 0$, padj < 0.05) across all three pairwise contrasts. (D) Three-way Venn diagram restricted to significantly downregulated DEGs ($\log_2 FC < 0$, padj < 0.05) across all three pairwise contrasts. (E) Heatmap of Z-score normalized FPKM expression values for the top 50 most significantly differentially expressed genes, displayed as individual biological replicates (all 9 samples as separate columns). (F) Heatmap of Z-score normalized FPKM expression values for significantly differentially expressed atherosclerosis-related genes in the LDLr$^{-/-}$+MPLs vs LDLr$^{-/-}$ contrast (padj < 0.05), displayed as individual biological replicates (9 samples).

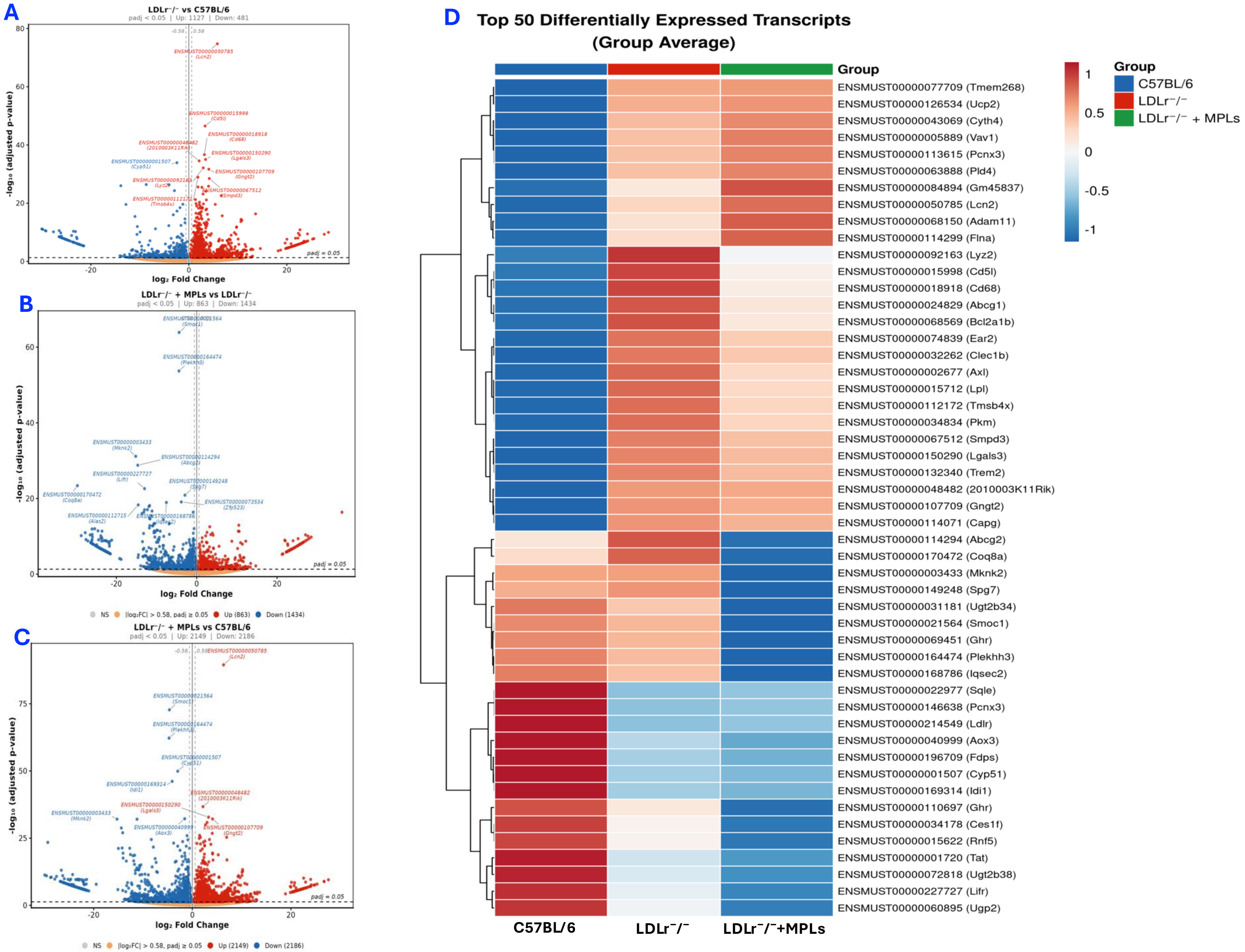


**Fig. S2. Transcript-level differential expression across contrasts reveals extensive isoform reprogramming in response to microplastic exposure.** (A–C) Volcano plots of transcript-level differential expression for (A) $LDLr^{-/-}$ versus C57BL/6, (B) $LDLr^{-/-}$+MPLs versus $LDLr^{-/-}$, and (C) $LDLr^{-/-}$+MPLs versus C57BL/6. Each point represents a transcript; the x-axis shows $\log_2$ fold change, and the y-axis shows $-\log_{10}$ adjusted p-value. Significantly upregulated transcripts (padj < 0.05, $\log_2FC > 0$) are shown in red and downregulated transcripts (padj < 0.05, $\log_2FC < 0$) in blue; non-significant transcripts are shown in grey. Dashed lines indicate significance (padj = 0.05) and fold-change (±0.58) thresholds. The top transcripts per contrast are annotated. (D) Heatmap of Z-score normalized expression (FPKM) for the top 50 most significantly differentially expressed transcripts (ranked across all contrasts), displayed as group averages.

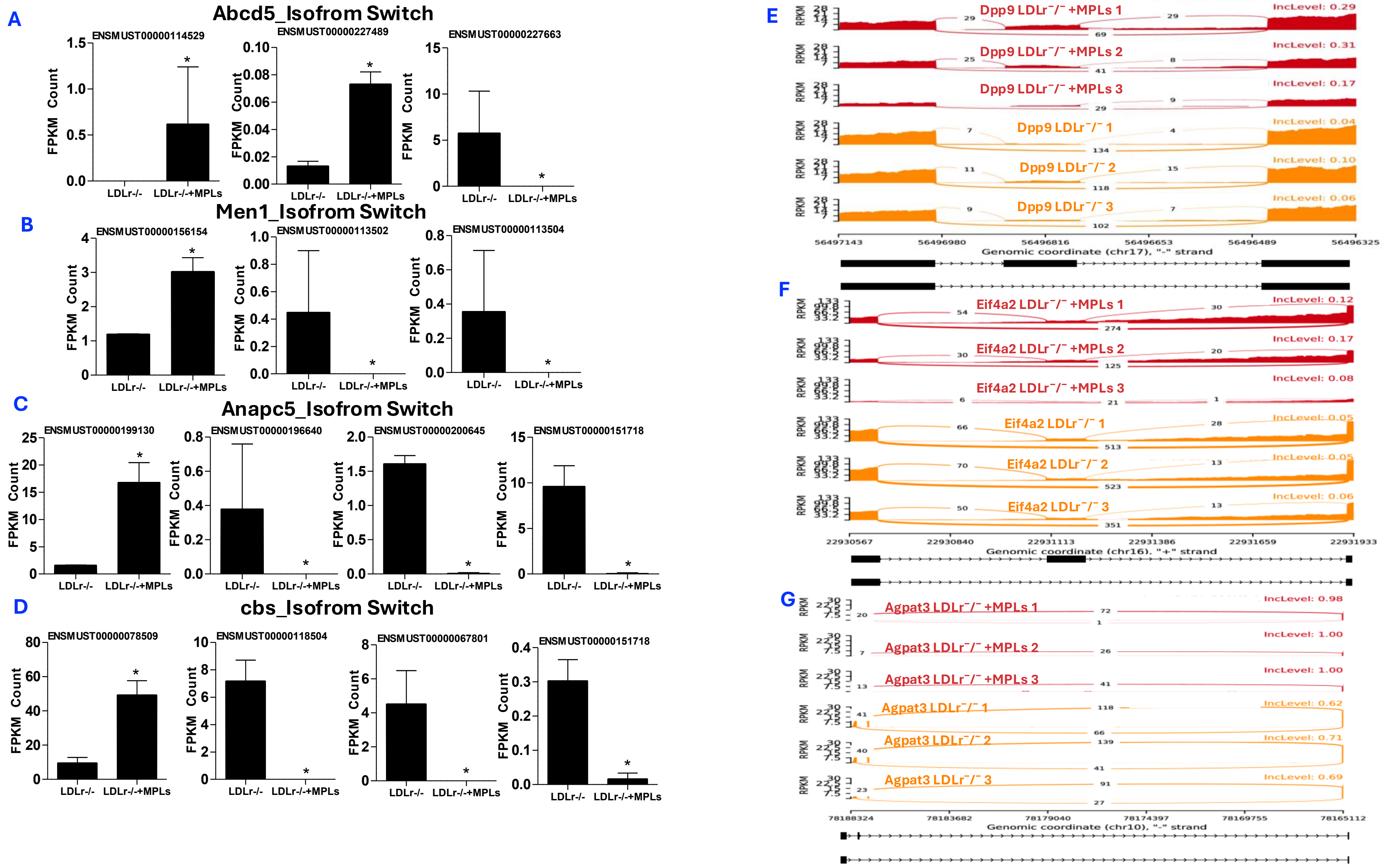


**Fig. S3. Additional isoform-switching events and read-level validation of alternative splicing in Nanoplastics-exposed LDLr⁻/⁻ liver.** (A–D) Representative isoform-switching events between LDLr⁻/⁻ and LDLr⁻/⁻+NPLs, showing mean transcript-isoform expression (FPKM ± SEM; padj < 0.05) for (A) Acbd5 (peroxisomal lipid transport), (B) Men1 (epigenetic regulation), (C) Anapc5 (cell-cycle control), and (D) Cbs (sulfur amino acid metabolism); in each, dominant isoforms are suppressed and minor isoforms induced. (E–G) Sashimi plots of representative rMATS alternative-splicing events (padj < 0.05) in LDLr⁻/⁻+NPLs (red) versus LDLr⁻/⁻ (orange), showing read coverage (RPKM) and junction counts per replicate, with shifts in exon inclusion for (E) Dpp9, (F) Eif4a2, and (G) Agpat3. All data are from liver, n = 3 per group, each replicate a pool of three mice.

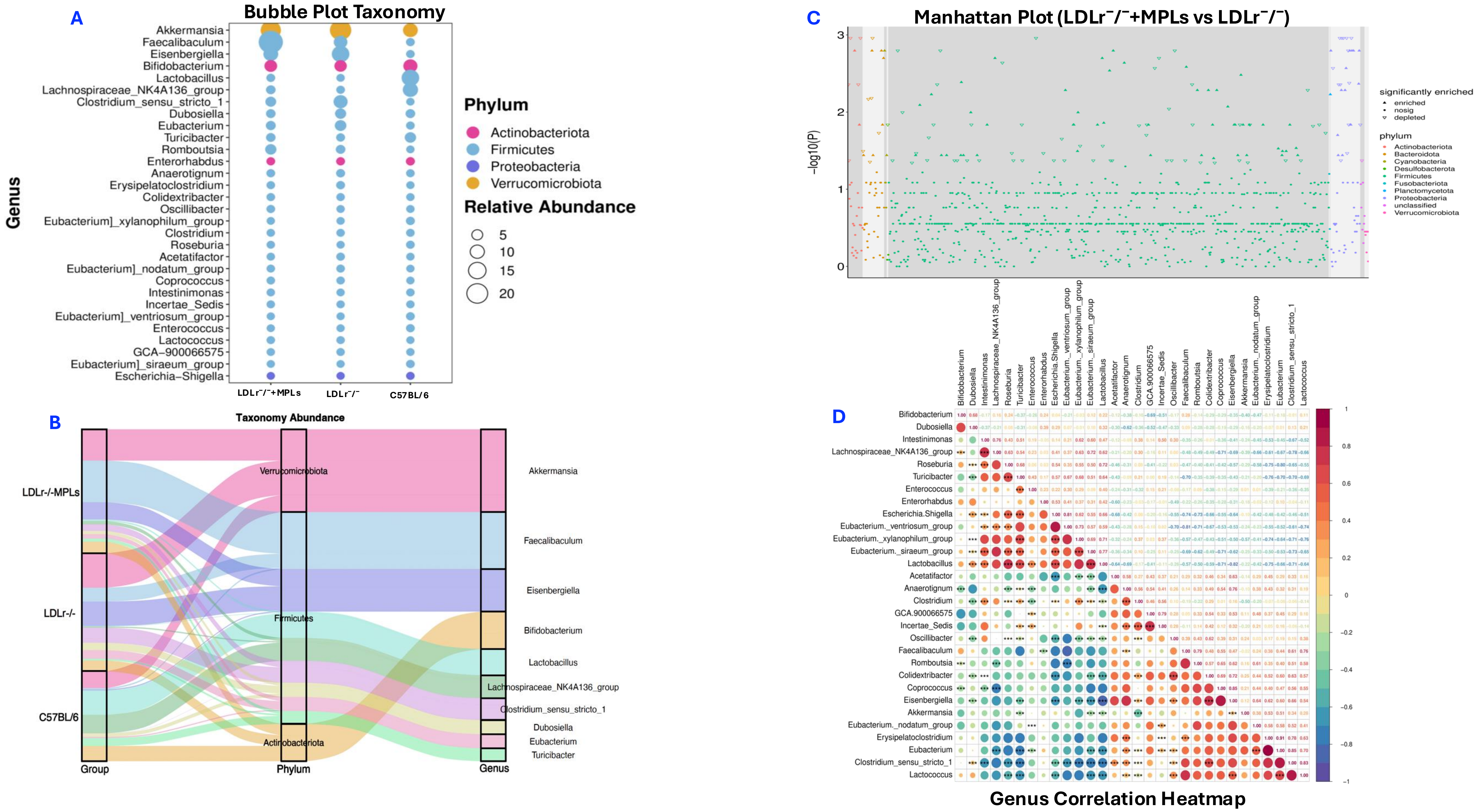


**Fig. S4. Genus-level microbial alterations and network restructuring in response to microplastic exposure.** (A) Indicator species analysis identifying group-specific genera, with *Faecalibaculum, Romboutsia, and Oscillibacter* showing highest indicator values. (B) Sankey plot illustrating microbial flow from groups to phylum and genus, highlighting enrichment of Faecalibaculum (LDLr⁻/⁻+MPLs), Eisenbergiella (LDLr⁻/⁻), and Lactobacillus (C57BL/6). (C) Manhattan plots showing significant differential abundance of genera across phyla ($-\log_{10}P \geq 2$), with broader shifts in LDLr⁻/⁻-MPLs vs LDLr⁻/⁻ (C) Correlation matrix revealing disrupted co-occurrence networks in LDLr⁻/⁻+MPLs, characterized by altered Firmicutes interactions and increased antagonistic relationships, indicating coordinated microbial dysbiosis.

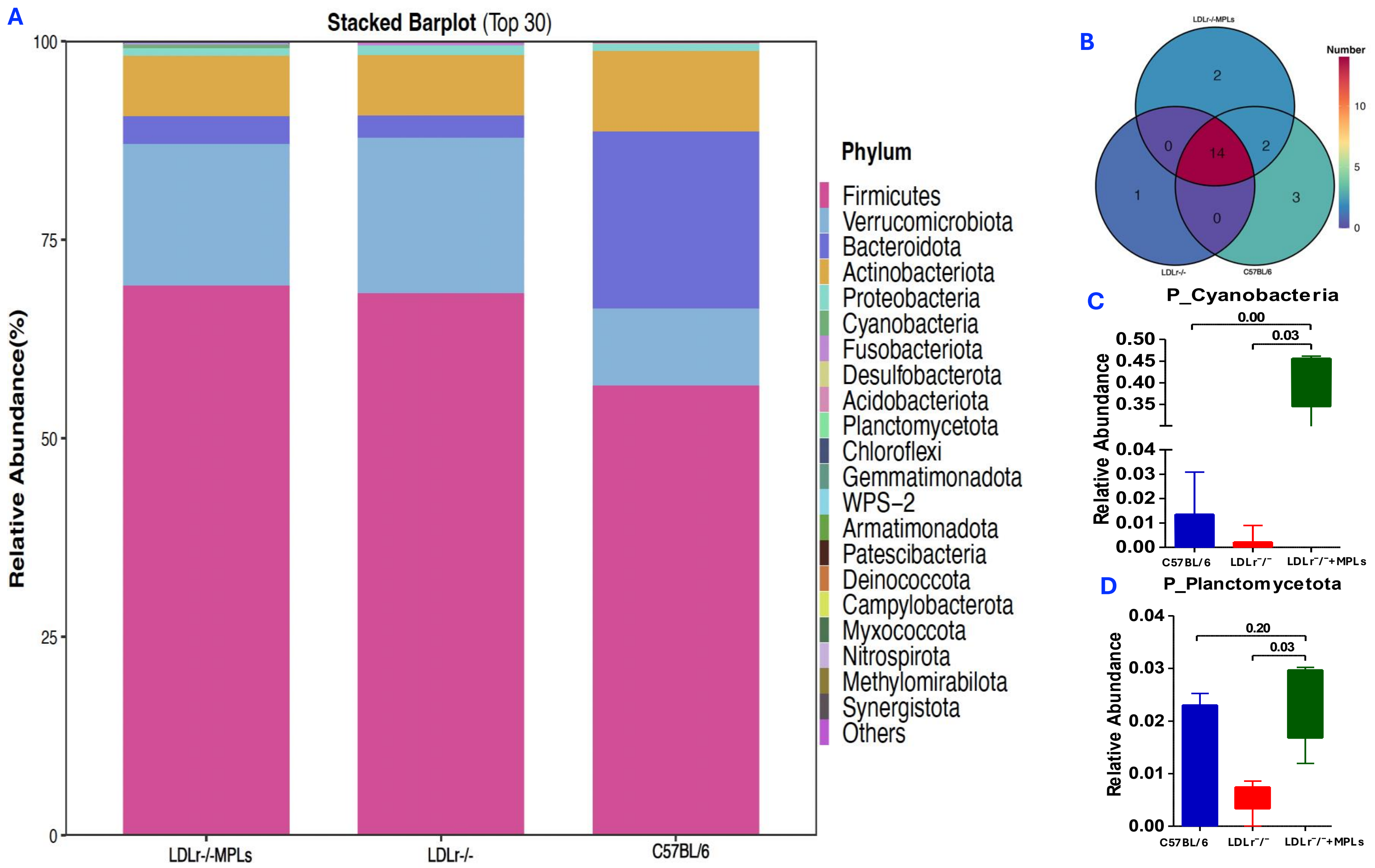


**Fig. S5. Nanoplastics induces Phylum-level microbial shift in $LDLr^{-/-}$ mice.** (A) Stacked bar plot showing the relative abundance of the top 30 bacterial phyla across $LDLr^{-/-}$ mouse + MPLs, $LDLr^{-/-}$, and C57BL/6 mouse groups. Microplastic exposure resulted in a clear compositional shift compared with $LDLr^{-/-}$ controls, indicating alterations in overall microbial community structure. (B) Venn diagram depicting shared and unique phyla among the three groups. Two phyla were uniquely detected in the $LDLr^{-/-}$+MPLs group, suggesting microplastic-associated microbial signatures. (C-D) Differential abundance analysis at the phylum level between $LDLr^{-/-}$+MPLs and $LDLr^{-/-}$ groups identified significant ($q < 0.05$) enrichment of *Planctomycetota* and *Cyanobacteria* following microplastic exposure, whereas no phyla were significantly depleted relative to $LDLr^{-/-}$ controls.

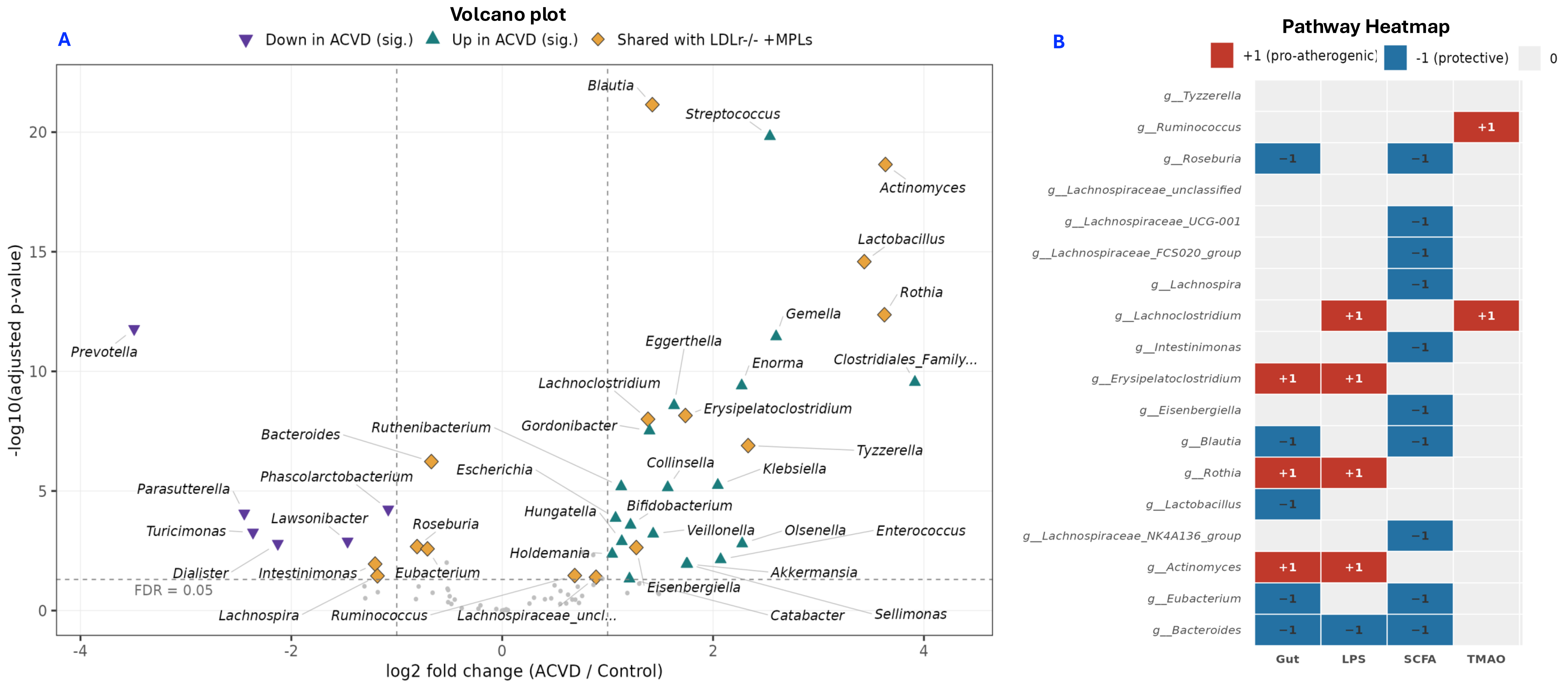


**Fig. S6. Human atherosclerosis-associated gut dysbiosis overlaps with the Nanoplastics-induced gut profile.** (A) Volcano plot of differential abundance results for human gut genera in atherosclerotic cardiovascular disease (ACVD) versus healthy controls (Jie 2017 cohort; n = 218 ACVD, n = 187 controls). The x-axis shows log2 fold change (ACVD/Control). The y-axis shows −log10(adjusted p-value). The horizontal dashed line marks the FDR threshold (padj = 0.05). Vertical dashed lines mark the ±0.5 log2FC effect size boundaries. Gold diamonds = 18 genera shared with the LDLr$^{-/-}$ +MPLs mouse signature (all padj ≤ 0.05; point size 4.8), teal upward triangles = genera significantly up-regulated in ACVD (log2FC > 0.5, padj ≤ 0.05), violet downward triangles = genera significantly down-regulated in ACVD (log2FC < −0.5, padj ≤ 0.05). (B) Heatmap showing mechanistic pathways implicated in Nanoplastics driver dysbiosis. Tile colors indicate pro-atherogenic, protective, or unestablished evidence. SCFA/butyrate depletion was the main pathway identified, followed by gut barrier disruption, LPS/endotoxin inflammation, and TMAO production.
