## Supplementary material for "Chronic Polystyrene Nanoplastics Exposure Reprograms Gene Expression, Alternative Splicing, and Disrupts Host–Microbiome–Metabolic Networks to Promote Atherosclerosis in LDLr⁻/⁻ Mice": https://uncg-my.sharepoint.com/:u:/g/personal/a_khan10_uncg_edu/IQDdOD-nqdMjS5LEbGvLDjUjAZGJiuZ5SVDtwyL_oY1JXp8?e=PeO5hi: Supplemantary Tables_S4_S5_S6_S7.docx

**Table 4.** MZmine 4.8.30 Peak Picking Parameters

| **Module** | **Parameters Used** |
| --- | --- |
| **A. Mass Detection (first, for MS1 data)** |  |
| 1. MS Level Filter 2. Polarity 3. Spectrum Type 4. Mass Detector 5. Noise Level | MS1, level 1  positive and negative  (processed in separate batches)  centroid  centroid  5.0E3 |
| **B. Mass Detection (second, for MS2 data)** |  |
| 1. MS Level Filter 2. Polarity 3. Spectrum Type 4. Mass Detector 5. Noise Level | MS2, level 2  positive and negative  (processed in separate batches)  centroid  centroid  100 |
| **C. Chromatogram Builder** |  |
| 1. MS Level Filter 2. Polarity 3. Spectrum Type 4. Min. Consecutive Scans 5. Min. Intensity for Consecutive Scans 6. Min. Absolute Height 7. *m/z* Tolerance (scan-to-scan) | MS1, level 1  positive and negative  (processed in separate batches)  centroid  5  5.0E3  1.0E5  0.003 *m/z* or 5.0 ppm |
| **D. Local Minimum Feature Resolver** |  |
| 1. MS1 to MS2 Precursor Tolerance (*m/z*) 2. Retention Time Filter 3. Minimum Relative Feature Height 4. Dimension 5. Chromatographic Threshold 6. Minimum Search Range RT 7. Minimum Absolute Height 8. Min. Ratio of Peak Top/Edge 9. Peak Duration Range 10. Minimum Scans (data points) | 0.003 m/z or 5.0 ppm  Use Tolerance, 0.20 min.  25%  Retention Time  70%  0.05  1.0E5  1.50  0.00 to 5.00  5 |
| **E. ^13^C Isotope Filter** |  |
| 1. *m/z* Tolerance (intra-sample) 2. Retention Time Tolerance 3. Monotonic Shape 4. Maximum Charge 5. Representative Isotope 6. Never Remove Feature with MS2 | 0.0015 *m/z* or 0.0 ppm  0.05 min  Checked  2  Most Intense  Checked |
| **F. Isotopic Peaks Finder** |  |
| 1. Chemical Elements 2. *m/z* Tolerance (feature-to-scan) 3. Maximum Charge of Isotope *m/z* 4. Search in Scans | H, C, N, O, S, Cl, Br, P, F  0.002 *m/z* or 0.0 ppm  2  Single Most Intense |
| **G. Join Aligner** |  |
| 1. *m/z* Tolerance (sample-to-sample) 2. Weight for *m/z* 3. Retention Time Tolerance 4. Weight for RT 5. Mobility Weight | 0.003 *m/z* or 5.0 ppm  2  0.10 minutes  1  1.0 |
| **H. Peak Finder (Gap Filling)** |  |
| 1. Intensity Tolerance 2. *m/z* Tolerance (sample-to-sample) 3. Retention Time Tolerance 4. Minimum scans | 20.0%  0.003 *m/z* or 5.0 ppm  0.10 minutes  5 |
| **I. Duplicate Peak Filter** |  |
| 1. Filter Mode 2. *m/z* Tolerance 3. RT Tolerance | New Average  0.0015 *m/z* or 0.0 ppm  0.05 minutes |
| **J. Feature Filter** |  |
| 1. Height 2. No. of Data Points | 1.0E5 to 5.0E10  5 to 500 |
| **K. Correlation Grouping (metaCorrelate)** |  |
| 1. RT Tolerance 2. Minimum Feature Height 3. Intensity Threshold for Correlation 4. Min. Samples Filter    a. Min Samples in All    b. Min Samples in Group    c. Min %-Intensity Overlap    d. Exclude Gap-Filled Features 5. Feature Shape Correlation    a. Min Data Points    b. Min Data Points on Edge    c. Measure    d. Min Feature Shape Correlation | 0.10 min  0.0E0  0.00  Max of 1 sample or 0.0%  Max of 0 sample or 0.0%  60.0%  Checked  5  2  Pearson  85% |
| **L. Ion Identity Networking** |  |
| 1. *m/z* Tolerance (intra-sample) 2. Check 3. Minimum Height 4. Ion Identity Library    a. Maximum Charge    b. Maximum Molecules/Cluster    c. Selected Adducts (positive)    d. Selected Adducts (negative) 5. Annotation Refinement    a. Delete Smaller Networks: Link Threshold | 0.0015 *m/z* or 0.0 ppm  One Feature  0.0E0  2  2  [M+H]^+^, [M+NH_4_]^+^, [M+Na]^+^, [M+K]^+^  [M-H]^-^, [M+Cl]^-^, [M+FA]^-^, [M+Acetate]^-^, [M+Br]^-^  4 |

**Table S5**. xMSannotator annotation parameters (A) and post-annotation filtering pipeline (B).

| **Parameter / Step** | **Value / Criterion** | **Description** |
| --- | --- | --- |
| ***A. xMSannotator Annotation Parameters*** | | |
| *Mass accuracy tolerance* | ±10 ppm | Maximum permitted deviation between observed and theoretical monoisotopic m/z for HMDB matching |
| *Retention time window* | 10 min | Window within which co-detected features are evaluated for adduct and isotope relationships |
| *Adduct/isotope correlation threshold* | r ≥ 0.70 | Minimum Pearson correlation required to confirm an adduct or isotope co-detection relationship |
| *Isotope peaks searched* | 5 | Maximum number of isotope peaks searched per feature |
| *Mass defect window* | 0.01 Da | Tolerance applied during isotope peak identification |
| *Reference database* | HMDB (union) | Human Metabolome Database; union strategy draws from all sub-databases to maximize coverage |
| *Ionization mode* | Positive (ESI+) | Consistent with C18 reversed-phase LC-MS data acquisition |
| *Primary anchoring adducts* | M+H | Primary positive-mode adduct used to anchor the annotation confidence score |
| *Adducts searched* | 15: M+H, M+NH4, M+Na, M+2H, M+H+NH4, M+ACN+H, M+ACN+Na, M+2ACN+H, 2M+H, 2M+Na, 2M+ACN+H, M+2Na−H, M+H−H2O, M+H−2H2O, M+2ACN+2H | Full set of positive-mode adduct forms searched during database matching |
| *NOPS elemental filter* | Enabled | Enforces chemically reasonable N–O–P–S atomic ratios to reduce false-positive annotations |
| *Confidence score threshold* | ≥ 2 | Only annotations supported by ≥2 independent evidence layers retained for downstream analysis |
| ***B. Post-Annotation Filtering Pipeline*** | | |
| **Filter Step** | **Criterion** | **Features Retained** |
| Raw xMSannotator output | — | 80,622 feature-annotation pairs (all candidates prior to filtering) |
| Confidence score filter | Score ≥ 2 | 11,990 pairs; low-confidence mass-match-only annotations removed |
| Primary adduct filter | M+H only | 8,705 non-redundant features; alternative adduct forms collapsed |
| Differential abundance | p < 0.05, Welch’s t-test (LDLr⁻/⁻+MPLs vs LDLr−/−) | 27 significant feature-annotation pairs |
| Deduplication by m/z–RT | Highest-confidence annotation per unique peak | unique confirmed differentially abundant features |

**Table S6**. Alpha diversity indices show within the sample diversity across group.

| **Analysis** | **group1** | **group2** | **p.adj** | **Significance** |
| --- | --- | --- | --- | --- |
| observed_species | LDLr⁻/⁻+MPLs | LDLr⁻/⁻ | 0.54 | ns |
| observed_species | LDLr⁻/⁻+MPLs | C57BL/6 | 0.54 | ns |
| observed_species | LDLr⁻/⁻ | C57BL/6 | 0.70 | ns |
| shannon | LDLr⁻/⁻+MPLs | LDLr⁻/⁻ | 0.36 | ns |
| shannon | LDLr⁻/⁻+MPLs | C57BL/6 | 0.07 | * |
| shannon | LDLr⁻/⁻ | C57BL/6 | 0.82 | ns |
| simpson | LDLr⁻/⁻+MPLs | LDLr⁻/⁻ | 0.36 | ns |
| simpson | LDLr⁻/⁻+MPLs | C57BL/6 | 0.04 | * |
| simpson | LDLr⁻/⁻ | C57BL/6 | 0.70 | ns |
| chao1 | LDLr⁻/⁻+MPLs | LDLr⁻/⁻ | 0.54 | ns |
| chao1 | LDLr⁻/⁻+MPLs | C57BL/6 | 0.59 | ns |
| chao1 | LDLr⁻/⁻ | C57BL/6 | 0.70 | ns |
| goods_coverage | LDLr⁻/⁻+MPLs | LDLr⁻/⁻ | 1 | ns |
| goods_coverage | LDLr⁻/⁻+MPLs | C57BL/6 | 0.43 | ns |
| goods_coverage | LDLr⁻/⁻ | C57BL/6 | 0.43 | ns |
| pielou_e | LDLr⁻/⁻+MPLs | LDLr⁻/⁻ | 0.20 | ns |
| pielou_e | LDLr⁻/⁻+MPLs | C57BL/6 | 0.04 | * |
| pielou_e | LDLr⁻/⁻ | C57BL/6 | 0.70 | ns |
| ace | LDLr⁻/⁻+MPLs | LDLr⁻/⁻ | 0.54 | ns |
| ace | LDLr⁻/⁻+MPLs | C57BL/6 | 0.59 | ns |
| ace | LDLr⁻/⁻ | C57BL/6 | 0.70 | ns |

**Table S7**. Adonis (PERMANOVA) results in group level differences in beta diversity.

| Term of interest: Group \| Permutations P-value: Pr(>F) | | | | | | |
| --- | --- | --- | --- | --- | --- | --- |
| Distance metric | Term | Df | SumOfSqs | R2 | F | Pr(>F) |
| Bray-Curtis | Group | 2 | 2.232364123 | 0.602933247 | 12.14774578 | 0.001 |
|  | Residual | 16 | 1.47014214 | 0.397066753 |  |  |
|  | Total | 18 | 3.702506264 | 1 |  |  |
| Jaccard | Group | 2 | 1.750292022 | 0.283958184 | 3.172531856 | 0.001 |
|  | Residual | 16 | 4.413615626 | 0.716041816 |  |  |
|  | Total | 18 | 6.163907648 | 1 |  |  |
| Unweighted UniFrac | Group | 2 | 0.963603354 | 0.338128286 | 4.086934424 | 0.001 |
|  | Residual | 16 | 1.886212508 | 0.661871714 |  |  |
|  | Total | 18 | 2.849815862 | 1 |  |  |
| Weighted UniFrac | Group | 2 | 0.7014656 | 0.578457909 | 10.97793878 | 0.001 |
|  | Residual | 16 | 0.511182009 | 0.421542091 |  |  |
|  | Total | 18 | 1.212647609 | 1 |  |  |
